# CPT1a regulates the delivery of extracellular fatty acids for cardiolipin turnover in prostate cancer cells

**DOI:** 10.1101/2024.01.16.575611

**Authors:** Nancy T Santiappillai, Mariam F Hakeem-Sanni, Anabel Withy, Lisa M Butler, Lake-Ee Quek, Andrew J Hoy

## Abstract

Mitochondrial fatty acid oxidation (FAO) has been proposed to be a major bioenergetic pathway in prostate cancer. However, this concept fails to consider FAO relative to other mitochondrial substrates. Here, we found extracellular long-chain fatty acids (LCFAs), including palmitate, stearate, oleate, linoleate, linolenate, are minor sources of carbon entering the TCA cycle compared to glucose and glutamine in prostate cancer cells, despite being assimilated in the mitochondria as acyl-carnitines. In contrast, cardiolipins were a prominent LCFAs sink, with some species achieving greater than 50% ^13^C-labelling within 6 hours, suggesting high cardiolipin turnover using extracellular LCFAs. Knockdown of CPT1a, the rate-limiting enzyme of LCFA entry into mitochondria, reduced the incorporation of extracellular linoleate into cardiolipins. These results demonstrate that FAO is not a major input for the TCA cycle and provide evidence for an underappreciated role for CPT1a in regulating LCFAs entry into mitochondria for cardiolipin remodelling.

## Introduction

Prostate cancer (PCa) display unique metabolic features compared to many other solid tumours, as it typically does not exhibit the “Warburg effect”. A hallmark of PCa is alterations in lipid metabolism, which arise from reports that PCa overexpress lipid metabolism genes^1–3^, have enhanced fatty acid (FA) oxidation (FAO) rates^4–6^, increased sensitivity to inhibition of FAO and FA synthesis pathways^7,8^, lipid profile alterations that associate with PCa progression^8^, and increased citrate oxidation for lipogenesis^9,10^ compared to benign epithelial cells and tissues. Of these features, it is the broader interpretations of the higher FAO rates that are poorly supported. In that, it has been long proposed that FAO is the predominant bioenergetic source in PCa^11,12^, yet there have been no quantifications of FAO rates in relation to other major mitochondria substrates, such as glucose and glutamine. PCa cells consume non-trivial amounts of glucose and glutamine^6,13^ and this shifts along a spectrum of glucose and lipid utilisation from benign to castration resistant PCa^14,15^. However, it is unclear whether FAO is a major carbon source for the TCA cycle, and thereby support the widely held belief that it is the predominant bioenergetic source, compared to glucose and glutamine in PCa cells.

Cytosolic LCFA-CoAs enter the mitochondria through the carnitine shuttle system^16^, with the contemporary view being that mitochondria LCFA-CoAs are shunted into β-oxidation to produce acetyl-CoA for the TCA cycle, and NADH and FADH_2_ for the electron transport chain (ETC)^17^. As such, CPT1 is viewed as the rate-limiting step of FAO, which is supported by evidence that loss-of-function decreases CO_2_ and ATP production and causes cell death in cancer cells^5,18–20^. This CPT1a-FAO-ATP-cell death cascade requires FAO to be the predominant bioenergetic process, and that glucose and glutamine metabolism are unable to sustain ATP levels in the setting of impaired FAO. Significant doubts about this cascade arose from Yao *et al*^21^ who reported that pharmacological inhibition of FAO with low concentrations of the CPT1 inhibitor etomoxir did not affect the proliferation rates of various cancer cells.

Secondly, they showed that CPT1a loss-of-function in BT549 breast cancer cells decreased mitochondrial lipid levels and altered mitochondrial morphology^21^. Similar observations have been reported in T cells^22^ and mouse heart^23^. Together, these results raise the possibility that intramitochondrial LCFA-CoAs are not a major TCA cycle source of carbons, and that CPT1 is essential for maintaining intramitochondrial lipid homeostasis and ultimately questions the sole commitment of mitochondria LCFA-CoAs into beta-oxidation. In this study, we aimed to determine whether FAO provides a significant carbon source to the TCA cycle in PCa cells and whether extracellular LCFAs contribute to other significant mitochondrial fates.

## Results

### Glucose and glutamine are the major TCA cycle carbon sources in prostate cancer cells

PCa is characterised by higher rates of FAO compared to prostate epithelial cells^4–6^, however, it remains to be determined how these rates relate to the use of other sources. We aimed to quantify, in parallel, the conversion of extracellular glucose, glutamine and LCFAs into TCA cycle metabolites in a panel of PCa cells spanning the spectrum of disease, from benign PNT1 cells to late-stage PC-3 cells. Consistent with previous reports^4–6^, PCa cells had 2-4-fold greater FAO rates compared to PNT1 benign cells (Fig 1a). These differences in baseline FAO rates could not be explained by active mitochondria amount (Fig. 1b, c), nor by CPT1a protein levels (Fig. 1d, e). Next, we performed parallel [U-^13^C] isotope tracing experiments to quantify the relative incorporation of uniformly labelled palmitate (FA 16:0), glucose, and glutamine carbons into TCA cycle metabolites. Palmitate was chosen as the representative LCFA as it is the most abundant FA circulating in humans^24^. We determined that 6 hours was the optimal endpoint for our tracing experiments as isotopic steady state was reached, and media substrates were not depleted (>50% of starting levels; Extended Data Fig. 1a). Similar studies using ^13^C-glucose and -glutamine in cultured cells have shown that 3 to 6 hours of label exposure is sufficient for isotopic steady state^25,26^. We separately noted that doubling of media volume to 2 mL was necessary for a 12 to 24 hour experiment, in order to maintain media substrates above 50% (Extended Data Fig. 1b). A notable pattern was the consumption of media lactate by cells that had depleted glucose (Extended Data Fig. 1a), thereby, demonstrating metabolic plasticity of PCa cells.

**Fig. 1:**
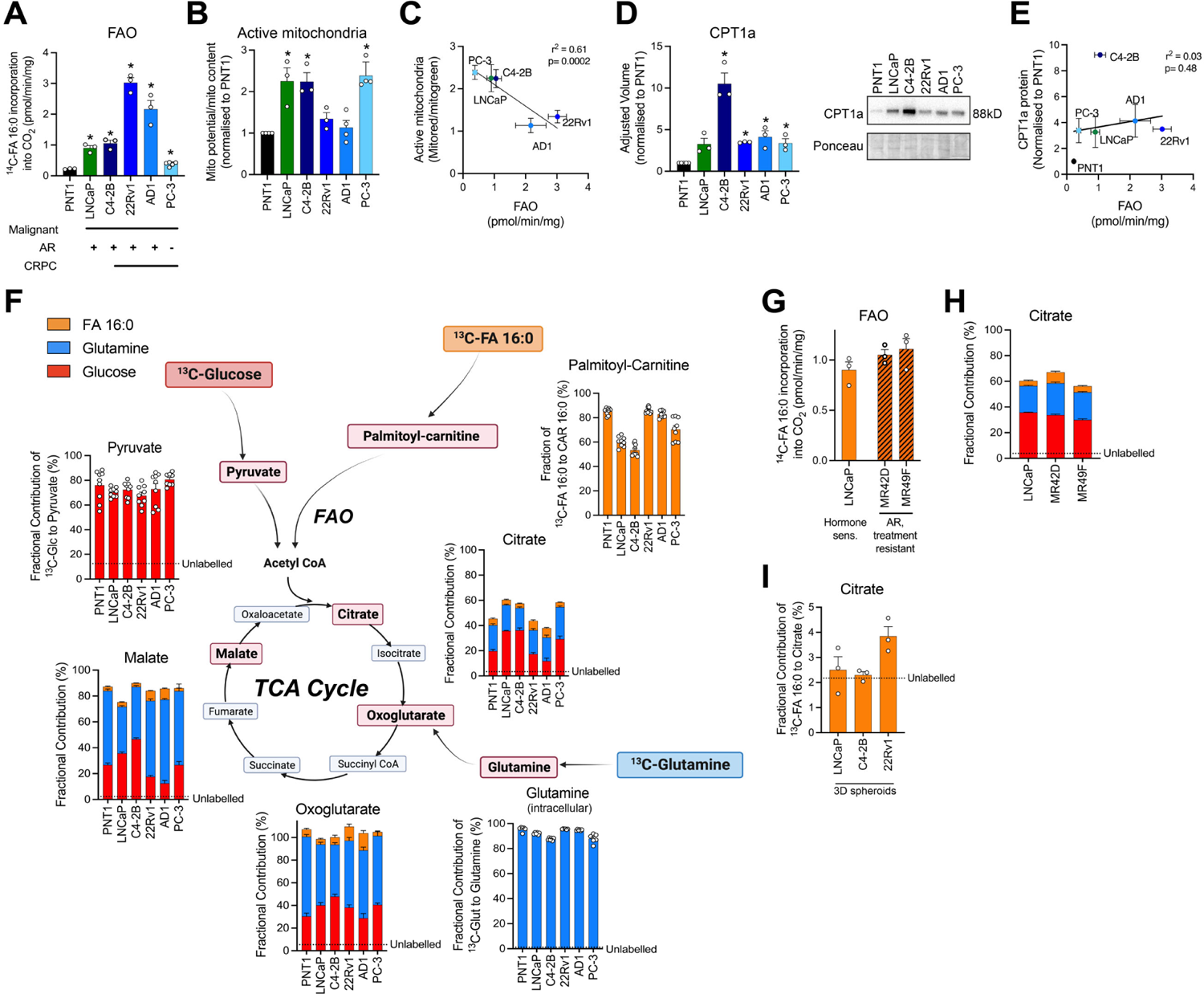
Palmitate is a minor TCA cycle substrate source compared to glucose and glutamine. (A) Fatty acid [1-^14^C]-FA 16:0 oxidation rates. N=3-5 per cell line. FAO rates were compared to PNT1 cells. (B) Active mitochondria as a fraction of mitochondrial potential to mitochondrial content normalised to PNT1 cells. N=3-4 per cell line. (C) Correlation of active mitochondria content to FAO excluding PNT1 cells. Error bars represent ± SD. P and r^2^ values by linear regression. (D) Quantification of CPT1a protein expression normalised to PNT1 cells. N=3 per cell line. A representative immunoblot is shown. (E) Correlation of CPT1a protein levels to FAO. Error bars represent ± SD. P and r^2^ values by linear regression. (F) Fractional contribution of [U-^13^C]-glucose to pyruvate (red), [U-^13^C]-glutamine to intracellular glutamine (blue), [U-^13^C]-FA 16:0 to palmitoyl-carnitine (orange) after 6 hours of labelling. [U-^13^C]-glucose, glutamine, or 16:0 to citrate, oxoglutarate and malate. N=9 per cell line. Level of natural enrichment of metabolite standards indicated. (G) Fatty acid [1-^14^C]-16:0 oxidation rates of MR42D and MR49F treatment resistant cell lines compared to treatment sensitive LNCaP cells. N=3 per cell line. (H) Fractional contribution of [U-^13^C]-glucose, glutamine, and FA 16:0 to citrate in MR42D, MR49F and LNCaP cells. N=9 per cell line. (I) Fractional contribution of [U-^13^C]-FA 16:0 to citrate in LNCaP, C4-2B and 22RV1 3D spheroid models. N=3 per cell line. Graphs show mean ± SEM unless stated otherwise. *p<0.05 by One-way ANOVA with Dunnett’s multiple comparisons test, unless stated otherwise.

We achieved sufficient labelling of the respective intracellular TCA cycle precursor pools after 6 hours (pyruvate >65%, glutamine >85%, and palmitoyl-carnitine >60%; Fig. 1f) and confirmed the uptake of extracellular labelled substrates. The fractional contribution of [U-^13^C]-glucose and [U-^13^C]-glutamine, calculated as the weighted sum by the number of labelled carbons^27^, combined ranged from 60 to 80% for all measured TCA cycle metabolites (Fig. 1f). In contrast, extracellular [U-^13^C]-palmitate contributed less than 15% of carbons to TCA cycle metabolites (Fig, 1f and Extended Data Fig. 1c-d). 22Rv1 and AD1 cells had the greatest enrichment of ^13^C-palmitate, which was consistent with their higher FAO rates (Extended Data Fig. 1c). In general, the fractional contribution of palmitate to TCA cycle metabolites weakly correlated with FAO and, somewhat surprisingly, inversely correlated with active mitochondria content (Extended Data Fig. 1d-e). Overall, non-trivial amounts of carbon from extracellular palmitate oxidation entered the TCA cycle, but this influx was dwarfed by the contribution of extracellular glucose and glutamine.

The treatment resistant PCa cells MR42D and MR49F^28^ have been reported to exhibit enhanced FA uptake and oxidation^29,30^ and so we next assessed the contribution of extracellular glucose, glutamine and palmitate to TCA cycle intermediates in these models. In our hands, FAO rates for both MR42D and MR49F cells were not different to treatment sensitive LNCaP cells (Fig. 1g). Likewise, the fractional contribution of palmitate to citrate in MR42D and MR49F cells was similar to LNCaP cells (4-8.5%), and consistent with our other results, citrate was largely derived from glucose (35%) and glutamine (20%; Fig. 1h and Extended Data Fig. 1g). We repeated our parallel tracing approach using 3D PCa spheroids as they exhibit a more oxidative phenotype compared to 2D cell cultures^31–33^. Again, we observed minimal contribution of palmitate into TCA cycle metabolites in LNCaP, C4-2B and 22RV1 3D spheroids (<6%; Fig. 1i and Extended Data Fig. 1j). Importantly, we confirmed that media glucose, glutamine, and palmitate were sufficiently plentiful prior to harvesting cells at 6 hours (Extended Data Fig. 1a, h), and that substantial ^13^C labelling of precursor pools were achieved in both treatment resistant cells and 3D spheroids (Extended Data Fig. 1f, i). Altogether, we show that extracellular glucose and glutamine are the major TCA cycle substrates compared to exogenous palmitate in a range of prostate cells, regardless of disease stage pathophysiology, treatment responsiveness, or multicellular microenvironment.

### Palmitate oxidation increases in the absence of glucose and glutamine but is insufficient to maintain TCA cycle function

One concern we had was that the low enrichment values (Fig. 1) were, in part, due to a technical or biological issue. As such, we set out to meaningfully increase the enrichment of extracellular palmitate into TCA cycle intermediates. Others have shown that limiting glucose availability increased FAO rates in MCF7 and T47D breast cancer cells^34^ and so we next determined whether FAO in PCa cells is increased by withdrawing glucose from the media, and hence lead to greater palmitate incorporation into the TCA cycle. Compared to basal FAO rates measured in the presence of 5 mM glucose, FAO in glucose-free conditions was 1- to 5-fold greater across the panel of prostate cells (Fig. 2a). We also measured FAO in a range of glucose concentrations (0 mM to 5 mM) and observed a dose-dependent suppression of FAO (Extended Data Fig. 2a). These data demonstrate that glucose levels, and glycolytic activity, impacts FAO rates.

**Fig. 2:**
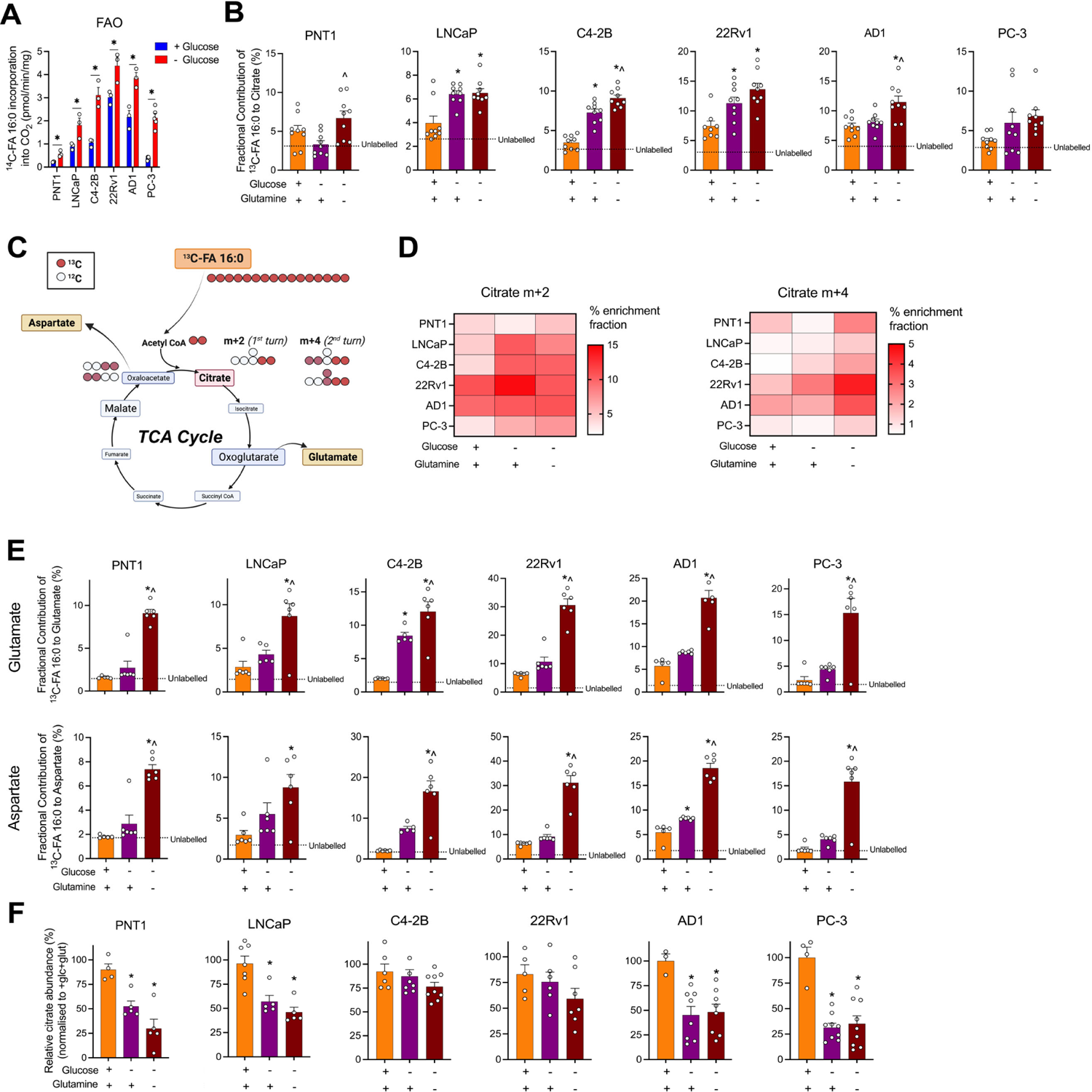
Palmitate alone is insufficient at maintaining TCA cycle activity. (A) [1-^14^C]-FA 16:0 oxidation rates in the presence (blue) or absence (red) of glucose. N=3-5 per cell line. Multiple unpaired t-tests. (B) Fractional contribution of [U-^13^C]-FA 16:0 to citrate with or without glucose or glucose and glutamine. N=9 per cell line. (C) Schematic tracing conversion of [U-^13^C]-FA 16:0 into citrate m+2 and m+4 isotopologues. (D) Enrichment fraction (%) of [U-^13^C]-FA 16:0 to m+2 and m+4 citrate isotopologues in +glucose/+glutamine, - glucose/+glutamine, and -glucose/-glutamine media conditions. N=6 per cell line. (E) Fractional contribution of [U-^13^C]-FA 16:0 to glutamate and aspartate +glucose/+glutamine, - glucose/+glutamine, or -glucose/-glutamine. N=5-6 per cell line. (F) Relative abundance (normalised to +glucose/+glutamine) of [U-^13^C]-FA 16:0 to citrate. N=4-9 per cell line. Graphs show mean ± SEM. P<0.05 by One-way ANOVA with Dunnett’s multiple comparisons test, unless stated otherwise. *compared to +glucose/+glutamine, ^compared to -glucose/+glutamine.

We next quantified the extent that extracellular palmitate provides carbons to TCA cycle metabolites in the absence of glucose or glucose and glutamine. There was no significant impact on cellular metabolic activity (MTT) or cell viability in glucose-free and glucose-/ glutamine-free media conditions at 6 hours (Extended Data Fig. 2b, c). Consistent with the increase in FAO rates (Fig. 2a), the fractional contribution of palmitate to citrate was increased in glucose-free and glucose-/ glutamine-free conditions, but it remained remarkably minimal (∼4% to ∼6-8%; Fig. 2b). Notably, PNT1 benign and AD1 PCa cells had increased palmitate labelling of citrate only in the absence of both glutamine and glucose, but not glucose alone (Fig. 2b), suggesting that these cells mobilised glutamine first instead of palmitate to compensate for glucose deprivation. In the absence of glucose and glutamine, palmitate produced markedly more m+2 and m+4 citrate isotopologues, which represent palmitate utilisation in the first and second turns of the TCA cycle, respectively (Fig. 2c, d). This suggests that palmitate carbons were retained in the TCA cycle for longer under glucose and glutamine deprivation. We also observed other dynamic changes in the usage of palmitate-derived carbons during glucose and glutamine deprivation, especially into cataplerosis-derived metabolites glutamate and aspartate (Fig. 2e), which are produced from oxoglutarate and oxaloacetate respectively (Fig. 2c). Firstly, we observed essentially no incorporation of ^13^C-palmitate into glutamate and aspartate in cells cultured in media containing physiological levels of glucose and glutamine (Fig. 2e). In glucose-free, and most strikingly in glucose-/glutamine-free conditions, we saw significant ^13^C-palmitate labelling into glutamate and aspartate (Fig. 2c, e). As aspartate synthesis is essential for cell proliferation^35^ and glutamate for glutathione synthesis and redox homeostasis^36^, our results suggest that prostate cells, in the absence of glucose or both glucose and glutamine, upregulate cataplerotic reactions in an attempt to maintain cellular function and viability. Together, these data demonstrate the dynamic adaptability of prostate cell palmitate metabolism to supply carbons to the TCA cycle when deprived of glucose only or glucose and glutamine.

FAO has been proposed to be the dominant bioenergetic source for PCa cells^11,12^. As such, we reasoned that FAO would be capable of sustaining TCA cycle activity in glucose or glucose-glutamine deprived conditions. In striking contrast to ^13^C-palmitate enrichment patterns, citrate, malate and oxoglutarate abundance were reduced in cells cultured in glucose-free and glucose- / glutamine-free settings for 6 hours compared to glucose and glutamine replete settings (Fig. 2f and Extended Data Fig. 2d). Specifically, citrate levels were decreased by 43-70% in most cells, except for C4-2B and 22Rv1 cells, which were not impacted by glucose- and glucose/glutamine-free conditions (Fig. 2f). We did observe striking reductions in malate and oxoglutarate levels (between 53-95%) in all cells in response to glucose and glutamine depletion (Extended Data Fig. 2d). Importantly, these results preceded the cell death which occurred between 6 and 24 hours of culturing (Extended Data Fig. 2b, c). Combined, these data show that even with the increased FAO rates to incorporate more palmitate carbon into the TCA cycle as well as into aspartate and glutamate, these changes were insufficient to maintain metabolite abundance and so TCA cycle activity that were followed by cell death.

### Extracellular LCFAs are minor TCA cycle substrates

Since palmitate is a minor carbon source for the TCA cycle (Fig. 1 and 2), we questioned whether this was a palmitate-specific phenotype. To address this, we repeated our experiments with a panel of non-essential (16:0, 18:0, 18:1) and essential (18:2, 18:3) LCFAs of varying saturation and carbon chain lengths. There was a similar trend of minimal contribution (2-18%) to TCA cycle metabolites observed in our panel of prostate cells, despite the diversity of LCFAs (Fig. 3a). 22Rv1 and AD1 cells had the greatest palmitate labelling and FAO rates amongst prostate cells (Fig. 1a and Extended Data Fig. 1c), and consistently exhibited greater utilisation of LCFAs by the TCA cycle compared to an unlabelled control (Fig. 3a). In general, there was no clear substrate preference between palmitate and other LCFAs to feed carbons to the TCA cycle (Fig. 1c), despite substantial labelling of respective acyl-carnitine species (0.8-1, ratio m^n^/m0+m^n^; Fig. 3b). Collectively, we show that in the presence of physiological glucose and glutamine, diverse extracellular LCFAs were not major carbon sources for the TCA cycle, despite substantial acyl-carnitine precursor labelling.

**Fig. 3:**
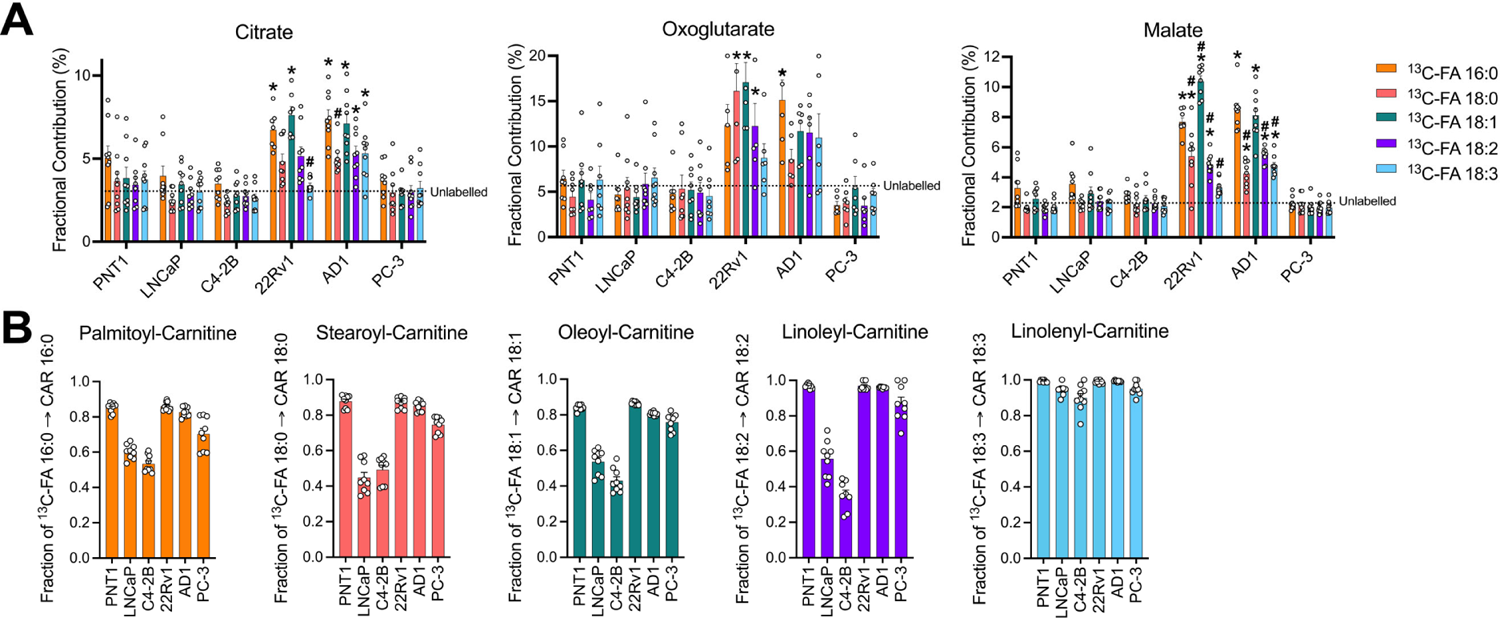
LCFAs of varying saturations are minor contributing substrate sources to the TCA cycle. (A) Fractional contribution of [U-^13^C]-FA 16:0, 18:0, 18:1, 18:2, 18:3 to citrate, oxoglutarate, malate. N=9 per cell line. (B) Fractional contribution of acyl-carnitines associated with the [U-^13^C]-FA substrates. N=9 per cell line. Graphs show mean ± SEM unless stated otherwise. P<0.05 by Two-way ANOVA with Tukey’s multiple comparisons test, *compared to unlabelled control (dotted line), #compared to [U-^13^C]-FA 16:0.

One possible reason for the disconnect between the substantial conversion of LCFA to acyl-carnitines and the lack of conversion to TCA cycle intermediates could be incomplete FAO, which generates medium chain acyl-carnitine species instead of fully oxidising LCFAs to acetyl-CoA^37,38^. In agreement with this hypothesis, FA 16:0, 18:0, and 18:1 greatly enriched shortened acyl-carnitine species (Extended Data Fig. 3). Specifically, FA 16:0 significantly enriched shortened CAR 14:0 (0.04-0.7 fraction) to CAR 10:0 (0-0.45 fraction), FA 18:0 enriched CAR 16:0 (0.01-0.38 fraction) to CAR 12:0 (0.04-0.4 fraction), and FA 18:1 enriched CAR 16:1 (0.03-0.79 fraction) to CAR 12:1 (0-0.1 fraction). The greatest amounts of exogenous FA enrichment into shortened acyl-carnitines were observed in PNT1, 22Rv1, AD1 and PC-3 cells (Extended Data Fig. 3), which were cells that also had the greatest incorporation of LCFAs into precursor acyl-carnitines (Fig. 3b). These data show that labelled extracellular LCFAs are not completely converted to acetyl-CoA, and thereby provides new insights into the relationships between acyl-carnitine synthesis, LCFA entry into mitochondria, and FAO.

### Extracellular LCFAs supports the rapid turnover of cardiolipin pools

Our observations showed that extracellular LCFAs were not completely oxidised to acetyl-CoA and minimally contributed carbons to the TCA cycle, despite being substantially present as acyl-carnitines (Fig. 3 and Extended Data Fig. 3). As such, we asked whether there are other pathways that could consume extracellular LCFAs after entering the mitochondria via CPT1a. This question arises from the work of Yao and colleagues^21^ as they were the first to report a reduction of mitochondrial complex lipids in BT549 breast cancer cells with CPT1a knocked down. Of their observed lipidome results, cardiolipins (CLs) piqued our interest as they are exclusively located within the inner mitochondrial membrane (IMM) and account for 10-20% of mitochondria phospholipid content^39^. CLs are composed of four fatty acyl chains of different saturation and chain lengths^40^ and provide essential structural functions to mitochondria membrane and cristae^41^, and ETC complexes I-IV for ATP production^42–44^. As CL remodelling functions to mature nascent CLs by the reacylation from donated acyl-CoAs^45^, it was conceivable that this process could be a significant consumer of mitochondria LCFA-CoAs. Therefore, we next investigated the contribution of extracellular LCFAs to CL synthesis and remodelling (Fig. 4a). First, we examined the enrichments of CL pools from extracellular [U-^13^C]-16:0, 18:0, 18:1, and 18:2 LCFAs. Surprisingly, cells cultured in media containing labelled LCFAs for 6 hours had >50% enrichment of many CL species, indicating substantial LCFA turnover of CL pools (Fig. 4b), in stark contrast to the marginal arrival of LCFA label into the TCA cycle (Fig. 1-3). CL has been shown to be a very slow turnover lipid class with CL half-life reported to be approximately two days in isolated rat cardiomyocytes^46^ and cultured rat H9c2 cardiomyoblast cells^47^. These turnover rates are much slower than other glycerophospholipids, including PC^46–49^. Here, we saw time-dependent enrichment of [U-^13^C]-FA 16:0 into PC 32:1 and 34:1 of C4-2B and PC-3 cells (Extended Data Fig. 4a) and that this enrichment was greater than related FA 16:0 containing CL species (CL 68:2 and CL 70:2) at 6 hours (Extended Data Fig. 4b). As such, C4-2B and PC-3 cells incorporate extracellular FAs into PCs at a higher proportion relative to CLs.

**Fig. 4:**
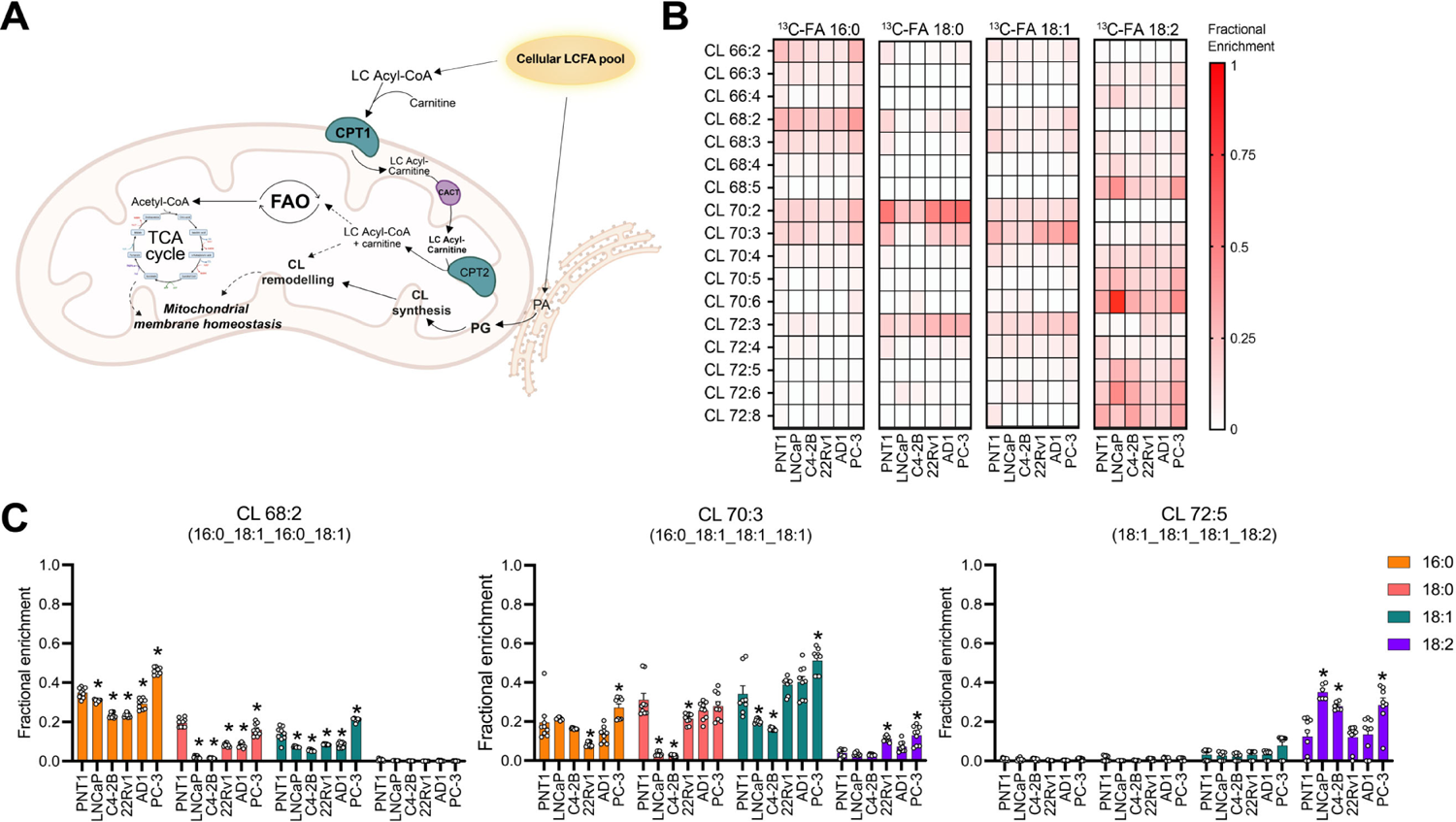
Extracellular LCFAs are directly and rapidly incorporated into cardiolipin pools. (A) Schematic depicting fates of FAs entering the mitochondria including FAO and CL pathways. (B) Fractional enrichment of [U-^13^C]-FA 16:0, 18:0, 18:1, 18:2 (150 μM) tracing to CL species for 6 hours. N=9 per cell line. (C) Fractional enrichment of [U-^13^C]-FA 16:0, 18:0, 18:1, 18:2 to CL 68:2, CL 70:3, CL 72:5 species. N=9 per cell line. Graphs show mean ± SEM. P<0.05 by One-way ANOVA with Tukey’s multiple comparisons test, *PCa cells vs PNT1 cells for each FA.

FAs, following activation into FA-CoAs, are substrates for elongation and desaturation reactions^50^. In our experiments, we saw that exogenous FAs 16:0, 18:0, and 18:2 were directly incorporated into the acyl-chains of CLs (i.e. unmodified via elongation or desaturation; Extended Data Fig. 4c). For instance, CL 68:2 is composed of 16:0 and 18:1 acyl-chains and was predominantly labelled by FA 16:0, while CL 70:3 is also composed of 16:0 and 18:1 acyl-chains but was mainly labelled by FA 18:1 (Fig. 4c). Similarly, CL 72:5 is composed of 18:1 and 18:2 chains and was almost entirely labelled by FA 18:2 within 6 hours (Fig. 4c). We observed some differences between PCa cells and benign PNT1 cells; however, no obvious pattern could be discerned (Fig. 4c). Finally, to compare the relative incorporation of the four FAs into CLs, we calculated an overall percentage of labelled fatty acyl chains among the 15 quantified CL species. In general, a greater proportion of FA 18:2 was labelled compared to FAs 16:0, 18:0, and 18:1 in our panel of prostate cells at 6 hours exposure to the respective substrates (Extended Data Fig. 4d). This suggests 18:2-containing CLs have the greatest turnover, which was also reported for cardiomyocytes and HeLa cells^51,52^. This may associate with the fact a majority of CL acyl chains is 18:2. The exceptions were LNCaP and C4-2B cells, which showed greater FA 16:0 13C enrichment among the CLs (32% and 14.2%, respectively) compared to FA 18:2 (9.5% and 8.2%, respectively; Extended Data Fig. 4d). Collectively, these data demonstrate that PNT1 prostate epithelial and a range of PCa cells rapidly incorporate extracellular LCFAs into CL acyl-chains.

### Reduced CPT1a protein levels decreased FAO and incorporation of LCFAs into CL

The high incorporation of extracellular LCFAs into CLs in both PCa cells and PNT1 cells was in stark contrast to the magnitude of enrichment into TCA cycle metabolites (Fig. 3 and 4). LCFAs are incorporated into CLs via multiple starting points, not exclusively competing with FAO for mitochondria FAs that are supplied via CPT1a. While these intramitochondrial acyl-CoAs are substrates for CL remodelling acyltransferases MLCAT1^45^ and ALCAT1^53^, CLs are also assembled by the *de novo* pathway producing nascent CLs from phosphatidylglycerol (PG)^54^, and then remodelled via acyl-chain exchange with PC and phosphatidylethanolamine by the actions of Tafazzin^55^. To tease out the association between CPT1a and CL remodelling, we quantified the incorporation of extracellular FA 18:2 in cells with CPT1a knocked down, since 18:2 has been reported to be predominantly incorporated into CLs via remodelling pathways^45,51^ (Fig. 5a). We included FA 16:0 in our experiments as it can be incorporated via both *de novo* synthesis and remodelling^56,45^. Further, we used C4-2B cells for this experiment as they had the greatest level of CPT1a protein in our panel of prostate cells (Fig. 1d). As expected, CPT1a knockdown (CPT1a^KD^; reduced by 89%) in C4-2B cells (Fig. 5b) decreased palmitate oxidation (Fig. 5c) and the abundance of ^13^C-labelled 18:2 and 16:0 acyl-carnitines compared to scrambled non-targeting control cells (Fig. 5d). CPT1a^KD^ decreased FA 16:0 enrichment of citrate from 3.5% to 3% (delta 0.5%) but did not impact the contribution from FA 18:2 (Fig. 5e).

**Fig. 5:**
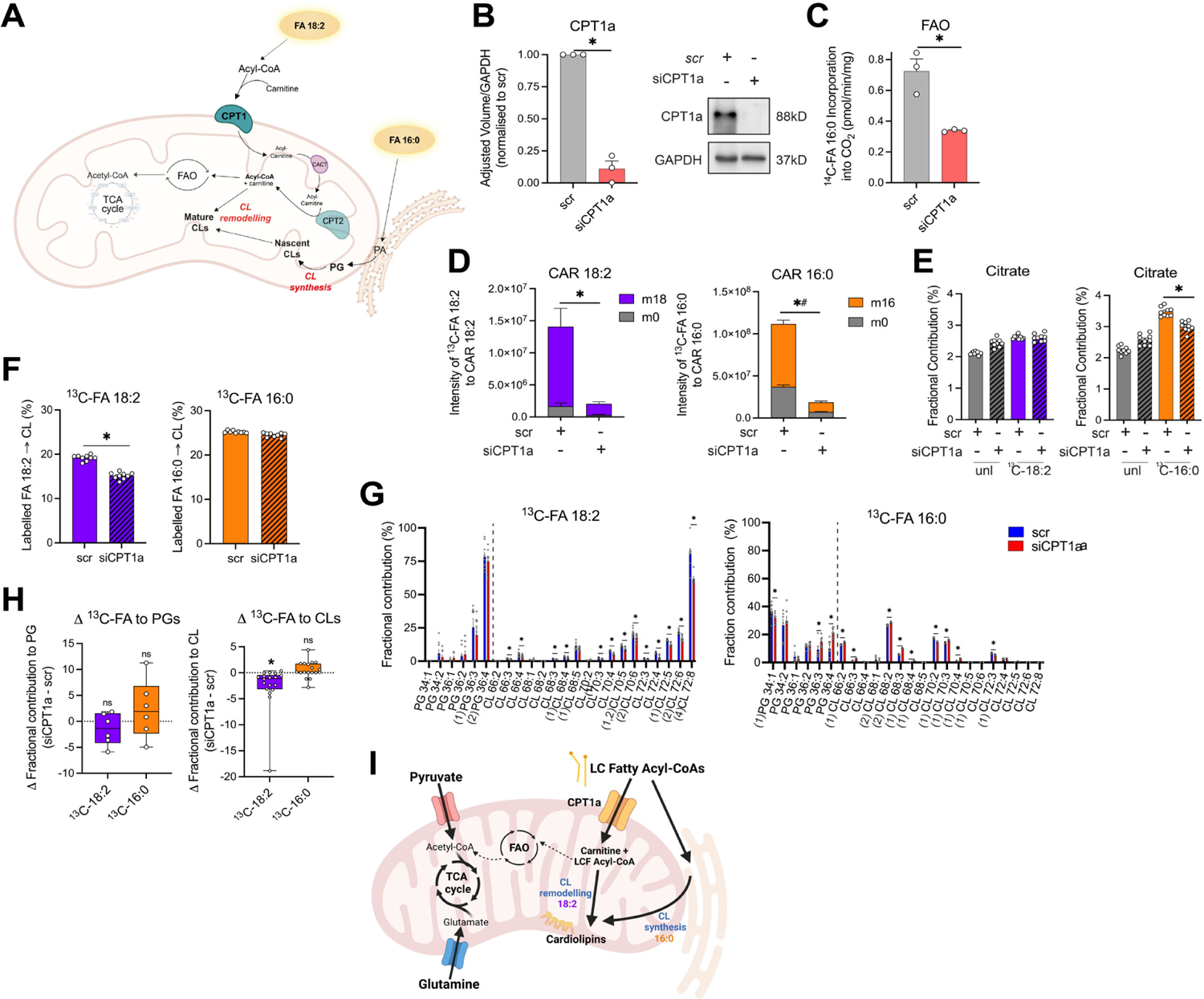
CPT1a knockdown reduced FA 18:2 derived CL remodelling. (A) Schematic depicting expected FA 18:2 to CL remodelling downstream of CPT1a and FA 16:0 to CL synthesis pathways derived from the endoplasmic reticulum. (B) CPT1a knockdown by reverse siRNA transfection in C4-2B cells for 72 hours quantified by immunoblot. Representative immunoblots shown. N=3 per group. (C) [1-^14^C]-FA 16:0 oxidation rates in non-targeting scrambled (scr) and siCPT1a C4-2B cells. N=3 per group. (D) Intensity of [U-^13^C]-FA 18:2 or 16:0 to carnitine species following siCPT1a treatment. N=9 per group. Statistically significant differences between labelled (m16 orange or m18 purple) intensities (*) or unlabelled (m0 grey) intensities (#). (E) Fractional contribution of unlabelled or [U-^13^C]-FA 18:2 or 16:0 to citrate in scr or siCPT1a C4-2B cells. N=9 per group. (F) Percentage of 18:2 or 16:0 acyl chain contained in CLs labelled by exogenous [U-^13^C]-FA. N=9 per group. (G) Fractional contribution of [U-^13^C]-FA 18:2 or 16:0 to PG and CL species. Annotated n values on the x-axis indicate the number of 18:2 or 16:0 containing acyl-chains. N=9 per group. Unpaired t-tests.(H) Delta differences between fractional contribution of [U-^13^C]-FA 18:2 or 16:0 to PG (N=6) and CL species (N=17). Statistically significant differences within [U-^13^C]-FA 18:2 or 16:0 groups to 0 difference determined by Nonparametric Wilcoxon signed rank test. Error bars represent ± min to max values. (I) Schematic depicting major carbon contributions from glucose and glutamine into the TCA cycle in comparison to the substantial contributions of LCFA-derived acyl-CoAs for CLs either by CL remodelling downstream of CPT1a or CL synthesis. Graphs show mean ± SEM unless stated otherwise. ns not significant, *p<0.05 between siCPT1a and scr control groups by student’s t-test unless stated otherwise.

In contrast, we found CPT1a^KD^ decreased FA 18:2 enrichment of CLs. The overall proportion of labelled FA 18:2 among the CL species was decreased from 19% to 15%, but this reduction was not reciprocated by FA 16:0 (Fig. 5f, Extended Data Fig. 5a). Namely, we saw all individual CL species had less FA 18:2 enrichment, but the effects were mixed for FA 16:0 enrichment (Fig. 5g). The latter suggests FA 16:0 is incorporated into CLs largely via the *de novo* pathway, which is independent of CPT1a and mitochondrial acyl-CoAs^54^. As expected, there was no difference in ^13^C-FA 18:2 and ^3^C-FA 16:0 PG enrichment between CPT1a^KD^ and scrambled control cells (Fig. 5h, Extended Data Fig. 5b, c), which suggests that the reduction in CPT1 activity did not divert LCFAs towards the endoplasmic reticulum to increase PG and CL synthesis. Altogether, we showed that that CPT1a regulates the incorporation of FA 18:2 into CL, thereby demonstrating that CL remodelling is a pathway downstream of CPT1a alongside FAO and the TCA cycle (Fig. 5i).

## Discussion

PCa do not exhibit classical Warburg metabolism but displays a more lipid dependent phenotype^8,57^. One aspect of this lipid phenotype that is widely accepted is the assumption that FAO is the predominant bioenergetic pathway in PCa, which likely arises from reports of enhanced FAO rates in PCa cells compared to benign cells^4–6^, low glycolytic activity at early disease stages^14,15^, and increased sensitivity to FAO targeting^7^. However, this belief that FAO is a dominant bioenergetic pathway in PCa, first proposed by Liu^11^ in 2006, failed to consider the contribution of other major respiration substrates. CPT1a has been implicated as the rate-limiting step of FAO because its inhibition results in decreased ATP production and cancer cell death^5,18–20^, and assumes that LCFAs are entirely diverted to FAO and the TCA cycle. Here, we generated evidence disputing this dogmatic view by showing extracellular LCFAs are minor carbon sources to the TCA cycle compared to glucose and glutamine in PCa cells, despite their increased FAO rates compared to benign PNT1 cells. We also showed that LCFAs are not completely oxidised to acetyl-CoA and produce shorter acyl-carnitine species. Most strikingly, there was substantial incorporation of LCFAs into CLs within 6 hours, and that CPT1a regulated the assimilation of FA 18:2 into CLs. Together, our data show a substantial role for CPT1a to regulate CL turnover in PCa cells and propose that observations of a CPT1a-dependent reduction of FAO and mitochondrial function can be explained, in part, by perturbed CL maturation and reduced oxidative phosphorylation efficiency.

We and others have shown that FAO in PCa cells is faster than benign prostate epithelial cells^4–6^, and that inhibiting CPT1 impacts cell viability^18–20^. These observations, combined with similar studies and the hypothesis article by Liu^11^, has resulted in the widely held view that FAO is a dominant bioenergetic pathway in PCa. Here, we present comprehensive evidence that challenges this. In a series of experiments, we show that extracellular LCFAs are a minor contributor of carbons to TCA cycle metabolites, compared to glucose and glutamine, which may, in part, be explained by incomplete oxidation. Further, FAO could not meet the anaplerotic demand to maintain TCA cycle activity in the absence of glucose alone or glucose and glutamine, leading to cell death. We also determined that extracellular FAs essentially do not contribute to glutamate and aspartate synthesis in the presence of extracellular glucose and glutamine, but do so in their absence. These results were seen across the spectrum of PCa progression, from benign to late-stage, 2D and 3D models. Importantly, the magnitude of exogenous FA enrichment into citrate was comparable to similar studies, including ∼15% in *ex vivo* cultured malignant Patient-Derived Xenografts (500 μM [U-^13^C]palmitate, 4 hours)^6^ and BT549 cells (100 μM [U-^13^C]palmitate, 24 hours)^21^, <15% in HCC1954 and SKBR3 breast cancer cells (100 μM [U-^13^C]palmitate, 24 hours)^58^, and ∼10% in MCF-7 and MDA-MB-231 breast cancer cells cultured in glucose-free media (100 μM [U-^13^C]palmitate, 4 hours)^59^. Therefore, these observations raise questions regarding the many studies demonstrating that CPT1 loss-of-function leads to cell death ^19,60^. One possible explanation is that many studies used etomoxir, an irreversible inhibitor of CPT1, at concentrations that have been reported to have significant off-target effects^21^, and often failed to identify the minimum concentration of etomoxir to maximally suppress FAO. Another reason could be due to the focus on demonstrating a dose-dependent reduction in cell viability and thereby determining IC_50_ in the absence of measures of FAO or CPT1 activity. Finally, the other technical consideration when comparing our results to others is the use of the Seahorse XF Analyser Palmitate Oxidation Kit, which measures oxygen consumption in saturating conditions of palmitate and carnitine but in media that is free of glucose, pyruvate, and glutamine. Further, the recommendation to culture cells in substrate limited conditions overnight will impact cellular metabolism. We observed striking increases in FAO rates, ranging from a doubling to a 5-fold increase, as well as substantial changes in intracellular partitioning of LCFA carbons in glucose-free conditions. As such, any observed changes in FAO performed in conditions free of major TCA cycle substrates must be interpreted with caution and likely explains the differences between our observations and other published studies. Unlike other studies, we showed that CPT1a^KD^ substantially decreased acyl-carnitine levels and the synthesis from extracellular LCFAs, and had a minor impact the contribution of LCFAs to TCA cycle metabolites in media containing 5 mM glucose and 1 mM glutamine. Overall, we conclude that extracellular LCFAs are not completely oxidised, and are a minor source of carbons for TCA cycle metabolites in a panel of PCa cells and benign PNT1 cells, therefore that FAO is not a major bioenergetic pathway.

In general, much of the literature discussing CPT1 function and mitochondrial FA metabolism state that CPT1 is rate-limiting for FAO and that FAO is the sole destination for intramitochondrial LCFA-CoAs following facilitated transport by the CPT1 – carnitine-acylcarnitine translocase – CPT2 system. This perception fails to consider that there are other destinations, especially given the evidence that CPT1 inhibition reduces mitochondrial lipid levels in cancer cells^21,61^ and alters mitochondrial morphology in BT549 cells^21^, T cells^22^, and rat heart muscle^23^. In our studies, we focused on the mitochondria exclusive CL as it is essential for mitochondria function and bioenergetics^62^. To our surprise, we saw rapid incorporation of extracellular LCFAs into CLs in PCa cells (>50% after 6 hours), which was much faster than what has been reported in isolate rat cardiomyocytes (2 days)^46^ and cultured rat H9c2 cardiomyoblast cells (∼54 hours)^47^. That said, we did determine that FA 18:2 was predominantly incorporated into the acyl-chains of CLs compared to other LCFAs, which is similar to previous reports^51,52^ and that the rate of incorporation of LCFAs into other lipid classes was faster than CL^46^. These results were especially important as cancer cells that substantially incorporate exogenous 18:2 into CL acyl-chains have enhanced stimulation of ETC complex I activity^51^ and reduced susceptibility to apoptosis^63^. These results showing significant enrichment of CLs with extracellular LCFAs provided a platform to determine whether CPT1 regulates LCFA incorporation into CLs, especially given that CPT1 loss-of-function reduces CL and mitochondrial lipids^21,61^. Consistent with our hypothesis, depletion of CPT1a reduced ^13^C-labelled FA 18:2 incorporation into CL in C4-2B PCa cells, and so suggest that CPT1 influences CL remodelling which may influence mitochondrial respiration and apoptosis^51,63^. Taken together, we propose that CLs, alongside FAO, are a prominent fate for extracellular LCFAs entering the mitochondria via CPT1a.

A limitation of our study is that we only quantified the contribution of extracellular substrates to mitochondrial pathways, which accounted for less than 65% of carbons to citrate. We focused on the relationship between FAO and TCA cycle metabolites, in part, to complement our previous reports that extracellular LCFAs as the major precursors for PCa lipids^5^. It is likely that the unaccounted for sources of carbons for TCA cycle metabolites include lactate^64^ or branched chain amino acids^65^, as well as intracellular sources of FAs and amino acids. Quantifying the totality could make new insights into carbon source for the TCA cycle and how differs in other cancer types, disease stage and impacted by the tumour microenvironment. We also provide evidence for incomplete FAO in PCa cells; however, further investigation into the roles of shortened acyl-carnitines may provide insights into other fates of LCFAs, alongside our new insights into CLs and TCA cycle biology. Finally, we showed for the first time that CPT1a impacts CL remodelling in PCa cells. Future experiments are required to link CPT1a function, CL homeostasis and ETC complex structure^62^ and other aspects of mitochondrial function, beyond FAO. This is especially true given that altered CL function has similar effects on mitochondrial morphology, function, and cell viability^66^ as CPT1a loss-of-function experiments^5,18–22^.

This study provides evidence that extracellular LCFAs are rapidly incorporated into CLs but are not the predominant carbon source to the TCA cycle in prostate cells compared to glucose and glutamine. Further, we conclude that CPT1a influences CL remodelling alongside FAO and so intramitochondrial LCFA-CoAs are not exclusively used for oxidation. Therefore, this work puts forward a new view of the role of CPT1 and its relationship to energy homeostasis and cell viability in PCa.

## Methods

### Key Resources Table

| REAGENT or RESOURCE | SOURCE | IDENTIFIER |
| --- | --- | --- |
| **Antibodies** |  |  |
| Mouse monoclonal anti-CPT1A | Abcam | Cat#Ab128568; RRID: AB 11141632 |
| Rabbit monoclonal anti-GAPDH | Cell Signalling Technology | Cat#2118; RRID: AB 561053 |
| **Chemicals, peptides, and recombinant proteins** |  |  |
| Enzalutamide MDV3100 | Selleckchem | Cat#S1250 |
| Lipofectamine RNAiMAX transfection reagent | Thermo Fisher Scientific | Cat#13778075 |
| [1-<sup>14</sup>C] Palmitic acid | Perkin Elmer | Cat#NEC075H250UC |
| Palmitic acid | Sigma-Aldrich | Cat#P0500 |
| D-Glucose | Sigma-Aldrich | Cat#273053 |
| L-Glutamine | Gibco | Cat#25030081 |
| Fatty acid free BSA | Bovogen | Cat#BSAS0.10 |
| L-carnitine hydrochloride | Sigma-Aldrich | Cat#C0283 |
| Mitotracker Green | Thermo Fisher Scientific | Cat#M7514 |
| Mitotracker Red | Thermo Fisher Scientific | Cat#M7512 |
| FCCP | Sigma-Aldrich | Cat#SML2959 |
| DTT | Sigma-Aldrich | Cat#D0632 |
| Protease and Phosphatase Inhibitor Cocktail, EDTA free | Astral Scientific | Cat#T-2496 |
| Thiazolyl blue tetrazolium bromide MTT reagent | Sigma-Aldrich | Cat#M2128 |
| Linoleic acid | Sigma Aldrich | Cat#L1012 |
| D-Glucose (U-<sup>13</sup>C<sub>6</sub>, 99%) | Cambridge Isotope Laboratories, Inc. | Cat#CLM-1396 |
| L-Glutamine (U-<sup>13</sup>C<sub>5</sub>, 99%) | Cambridge Isotope Laboratories, Inc. | Cat#CLM-1822-H |
| Palmitic acid (U-<sup>13</sup>C<sub>16</sub>, 98%) | Cambridge Isotope Laboratories, Inc. | Cat#CLM-409 |
| Stearic acid (U-<sup>13</sup>C<sub>18</sub>, 98%) | Cambridge Isotope Laboratories, Inc. | Cat#CLM-6990 |
| Linoleic acid (U-<sup>13</sup>C<sub>18</sub>, 98%) | Cambridge Isotope Laboratories, Inc. | Cat#CLM-10487 |
| Linolenic acid (U-<sup>13</sup>C<sub>18</sub>, 98%) | Cambridge Isotope Laboratories, Inc. | Cat#8386 |
| **Critical commercial assays** |  |  |
| Pierce BCA Protein assay kit | Thermo Fisher Scientific | Cat#23225 |
| Immobilon Crescendo Western HRP substrate | Merck Millipore | Cat#WBLUR0500 |
| CellCarrier Spheroid ULA 96-well microplates | Perkin Elmer | Cat#6055330 |
| **Experimental models: Cell lines** |  |  |
| PNT1 | ECACC | 95012614 |
| LNCaP | ATCC | CRL-1740 |
| C4-2B | ATCC | CRL-3315 |
| 22Rv1 | ATCC | CRL-2505 |
| CWR-R1-AD1 | A/Prof. Luke Selth, Flinders University | Nyquist 2013<sup>67</sup>, Li 2013<sup>68</sup> |
| PC-3 | ATCC | CRL-1435 |
| MR42D | Prof. Lisa Butler, SAHMRI | Bishop 2017<sup>28</sup> |
| MR49F | Prof. Lisa Butler, SAHMRI | Bishop 2017<sup>28</sup> |

| **Oligonucleotides** |  |  |
| --- | --- | --- |
| ON-TARGETplus Human CPT1A siRNA SMARTpool | Horizon Discovery | Cat#L-009749-00-0005 |
| ON-TARGETplus Non-targeting Control pool | Horizon Discovery | Cat# D-001810-10-05 |
| **Software and algorithms** |  |  |
| GraphPad Prism V9 | GraphPad Software | https://www.graphpad.com/ |
| Image Lab 6.1 | Bio-Rad | https://www.bio-rad.com/en-au/product/image-lab-software |
| FlowJo 2 | FlowJo LLC | https://www.flowjo.com/ |
| MATLAB | Mathworks | https://au.mathworks.com/products/matlab.html |
| MSConvert | N/A | Chambers 2012<sup>69</sup> |
| MSDial | Riken | http://prime.psc.riken.jp/compms/msdial/main.html |
| Biorender | Biorender | https://biorender.com/ |

### Method Details

#### Cell lines and culture conditions

Human benign prostate epithelial cell line PNT1 and human prostate carcinoma cells lines LNCaP (androgen receptor (AR) +, hormone sensitive), C4-2B, 22Rv1, AD1 (AR+, androgen independent), and PC-3 (AR-, hormone resistant) cells were cultured in RPMI1640 medium (Life Technologies), supplemented with 10% FBS (Cytiva Hyclone), and 1% penicillin/streptomycin (Gibco). MR42D and MR49F (hormone resistant, treatment resistant) cells were cultured in RPMI 1640 medium (Life Technologies), supplemented with 10% FBS, 1% penicillin/streptomycin, and 10μM enzalutamide. All cells were incubated at 37°C and 5% CO_2_.

#### Spheroid culturing

Cells were seeded in 96-well Spheroid microplates, at a density of 5 000 cells/well (LNCaP), 8 000 cells/well (22Rv1), and 2 000 cells/well (C4-2B), in RPMI medium in 100 μL per well. An additional 100 μL of medium was added to cells at 24h and left un-agitated for a further 24h until spheroids formed.

#### Cell Transfection

C4-2B cells were transfected for 72 hours using RNAiMAX transfection reagent and 25 pmol pooled CPT1a siRNA or non-targeting scrambled control, according to the manufacturer’s instructions. After 72 hours, cells were used for subsequent radiolabelling, western immunoblotting, and ^13^C-labelling experiments.

#### ^14^C-labelling

Cells were washed in warm PBS and incubated in 600 μL media containing 0.1 mmol/L cold palmitate, [1-^14^C]-palmitate (0.1μCi/mL) conjugated to 2% (wt/vol) FA-free BSA and 1 mmol/L -carnitine in RPMI glucose-free medium (Gibco) for 4 hours. Fatty acid oxidation rates were determined from ^14^CO_2_ production as previously described^1^ and supplemented with 0 mM or 5 mM glucose to measure metabolic flexibility. Medium for dose dependent glucose deprivation results were supplemented with 0 mM, 1 mM, 3 mM, 5 mM glucose. Protein content was measured by BCA protein assay and absorbances read at 540 nm using a TECAN infinite M1000 pro plate reader.

#### Mitotracker

Cells were seeded in 12-well plates at a density of 2×10^5^ cells/well in RPMI medium. After 24 hours, cells were washed in warm PBS and obtained by trypsinisation and collected in serum-free RPMI medium. The cell suspension was centrifuged at 1500 rpm for 5 min in fluorescence-activated cell sorting (FACS) tubes. Cells were subsequently stained with 200 nM Mitotracker Red and Mitotracker Green at 37°C for 30mins. Additionally, unstained, red only, green only, and 2.5 μM FCCP treated cells were included as experimental controls. Cells were centrifuged at 1500 rpm for 5min and fixed with 4% paraformaldehyde at 37°C for 10min. Cells were washed with warm PBS and centrifuged at 1500 rpm for 5min and resuspended in 150 μL PBS before the samples were read on a BD LSR Fortessa flow cytometer. Data analysis was performed using FlowJo software.

#### Immunoblotting

Cells were seeded in 6-well plates (5×10^5^). After 24 hours, cells were lysed in RIPA buffer containing 1% DTT and 0.1% protease and phosphatase inhibitor cocktail. Protein content was measured by BCA protein assay and absorbances read at 540 nm using a TECAN infinite M1000 pro plate reader. Cell lysates were subjected to SDS-PAGE, transferred to polyvinylidene difluoride (PVDF) membranes (Merck Millipore). Ponceau staining or GAPDH was used as a loading control, and then membranes were immunoblotted for CPT1a. Chemiluminescence was performed using Western HRP Substrate and imaged using the Bio-Rad ChemiDoc MP Imaging System (Bio-Rad Laboratories) and Image Lab software.

#### Cell viability

Cells were seeded in 96-well plates, in 6 well replicates (3×10^3^ cells/well). After 24 hours the media was removed and replaced with glucose- and glutamine-free media (Gibco) supplemented with 5 mM glucose, 1 mM glutamine and/or 150 μM palmitate, conjugated to 2% (wt/vol) FA-free BSA. For glucose-deprived conditions DMEM no glucose, no glutamine media was supplemented with 1mM glutamine and 150 μM palmitate; for glucose and glutamine-deprived conditions media was supplemented with 150 μM palmitate only. Media (5 mM glucose, 1 mM glutamine, 150 μM palmitate) supplemented with 10% FBS was used as a control. At 0, 6, 12 and 24 hours, 50 μL of MTT reagent was added to wells. MTT reactions were incubated at 37°C and 5% CO_2_ for 4 hours, and then stopped by replacing media with 100 μL DMSO (Sigma–Aldrich) per well. Absorbances were read at 540 nm using a TECAN infinite M1000 pro plate reader. For live cell counts, cells were seeded in 6-well plates at 5×10^5^ cells/well. After 24 hours, media was replaced as described for the MTT assay, and cells counted by Trypan blue (Gibco) exclusion at 6 and 24 hours.

#### U-^13^C stable isotope tracing

Cells were seeded in 6-well plates (5×10^5^) for 2D culture, or as above for 3D spheroid models. After 24 hours, wells were washed with warm PBS and media replaced with 600 μL of DMEM no glucose (0 mM), no glutamine (0 mM) media (Gibco), supplemented with 5 mM glucose, 1 mM glutamine and 150 μM FAs replaced with their respective [U-^13^C] forms (Palmitate (16:0), Oleate (18:0), Stearate (18:1), Linoleate (18:2), or Linolenate (18:3)). For glucose-deprived conditions, DMEM no glucose, no glutamine media was supplemented with 1mM glutamine and [U-^13^C]-150 μM palmitate; for glucose and glutamine-deprived conditions, media was supplemented with [U-^13^C]-150 μM palmitate only. FAs were conjugated to 2% (wt/vol) FA-free BSA containing media for 24 hours at 37°C prior to treating cells. Cells were incubated in ^13^C containing medium for 6 hours, or up to 48 hours for time course experiments. For spheroid experiments, 200 μL of spheroids was pooled and collected and centrifuged at 1400 rpm for 5 min at room temperature. The supernatant medium was collected and used for further extracellular substrate experiments. The remaining cell pellet was washed with 2 mL cold PBS and centrifuged again. The cell pellet was collected for subsequent LC-MS analysis.

#### Extracellular substrate sampling

Cells were seeded in 6-well plates (5×10^5^) for 2D culture of as above for 3D spheroid models. After 24 hours, wells were washed with warm PBS and media replaced with 1 mL or 2 mL of DMEM no glucose, no glutamine media (Gibco) supplemented with 5 mM glucose, 1 mM glutamine and 150 μM FAs. 100 μL of extracellular media were collected from wells at 3, 6, 12, 24 hour timepoints for cells incubated in 1 mL of media, and 12, 24 hours for cells incubated in 2mL of media. For spheroid experiments, ^13^C labelled medium was collected from cells at 0 and 6 hours timepoints. Media samples were centrifuged at 16 000xg for 5 min at 4°C and the supernatant collected for subsequent LC-MS analysis.

#### Cell lysate extraction for LC-MS

Media was aspirated from plates and the cells were washed with 2 mL ice-cold 0.9% (w/v) NaCl. Cells were scraped in 300 μL of extraction buffer, EB, (1:1 LC/MS methanol:water (Optima) + 0.1x internal standards (non-endogenous polar metabolites) and transferred to a 1.5 mL microcentrifuge tube. A further 300 μL of EB was added to the cells and combined in the tube. 600 μL of chloroform (Honeywell) was added before vortexing and incubating on ice for 10 min. Tubes were vortexed briefly and centrifuged at 15 000g for 10min at 4°C. The aqueous upper layer was collected and dried without heat, using a Savant SpeedVac (Thermo Fisher) for metabolomics LC-MS analysis. The lower layer was collected and dried under nitrogen flow for lipidomics LC-MS analysis. For time course lipidomics experiments, extractions were carried out as previously described^70^ by quenching with 0.1M formic acid, neutralising with ammonium bicarbonate, and using 40:40:20 acetonitrile:methanol:water with 0.1 M formic acid as the solvent system.

#### Measuring stable-isotope labelled metabolites by LC-MS

Dried aqueous upper phase samples were resuspended in 40 μL Amide buffer A (20 mM ammonium acetate, 20 mM ammonium hydroxide, 95:5 HPLC H_2_O: Acetonitrile (v/v)) and vortexed and centrifuged at 15, 000g for 5min at 4°C. 20μL of supernatant was transferred to HPLC vials containing 20 μL acetonitrile for LCMS analysis of amino acids and glutamine metabolites. The remaining 20 μL of resuspended sample was transferred to HPLC vials containing 20 μL LC-MS H_2_O for LCMS analysis of glycolytic, pentose phosphate pathway (PPP), and TCA cycle metabolites. Amino acids and glutamine metabolites were measured using Vanquish-TSQ Altis (Thermo) LC-MS/MS system. Analyte separation was achieved using a Poroshell 120 HILIC-Z Column (2.1×150 mm, 2.7 μm) (Agilent) at ambient temperature. The pair of buffers used were Amide buffer A and 100% acetonitrile (Buffer B), flowed at 200 μL/min; injection volume of 5 μL. Glycolytic, PPP and TCA cycle metabolites were measured using 1260 Infinity (Agilent)-QTRAP6500+ (AB Sciex) LC-MS/MS system. Analyte separation was achieved using a Synergi 2.5 μm Hydro-RP 100A LC Column (100×2 mm) at ambient temperature. The pair of buffers used were 95:5 (v/v) water:acetonitrile containing 10 mM tributylamine and 15 mM acetic acid (Buffer A) and 100% acetonitrile (Buffer B), flowed at 200 μL/min; injection volume of 5 μL. Raw data from both LC-MS/MS systems were extracted using MSConvert^69^ and in-house MATLAB scripts.

#### Measuring extracellular substrates by LC-MS

20 μL of collected extracellular media was mixed with 80 μL water, vortexed. 10 μL of diluted media was mixed with 90 μL of extraction buffer containing 1:1 (v/v) acetonitrile and methanol + 1x internal standards (non-endogenous standards) at −30°C. The mixture was centrifuged at 12 000xg for 5 min at 4°C and transferred into HPLC vials for LC-MS analysis measured using the Vanquish-TSQ Altis as described above. Raw data from both LC-MS/MS systems were extracted using MSConvert^69^ and in-house MATLAB scripts.

#### Measuring stable isotope labelled lipids by LC-MS

Dried lower phase samples were resuspended in 100μL of 4:2:1 (v/v) isopropanol: methanol: chloroform containing 7.5 mM ammonium formate. Cardiolipin and acyl-carnitine lipids were measured using a Thermo Scientific Q-Exactive-HF-X Hybrid Quadrupole Orbitrap LC-MS/MS system.

For cardiolipins, analyte separation was achieved using an Agilent Poroshell 120, EC-C18, 2.1×150mm, 2.7μm column. The pair of buffers used were 60:40 acetonitrile: water (v/v) with 10 mM ammonium formate and 0.1% formic acid (mobile phase A), and 90:10 isopropanol: acetonitrile (v/v) with 10 mM ammonium formate and 0.1% formic acid (mobile phase B), flowed at 200 μL/min on negative mode. MS1 data was acquired with the following settings: 3.5kV, capillarity temperature 300°C, 120 000, injection time 100 ms, AGC 1×10^6^, scan range 1100-1650. For ddMS2 data acquirement, the following settings were used: top 20, resolution 30,000, 200-2000, isolation 1.0m/z, nce 30, AGC target 1×10^3^, intensity threshold 5×10^3^, dynamic exclusion 20 s. Raw data from both LC-MS/MS systems were extracted using MSConvert^69^, in-house MATLAB scripts, and MSDial.

For acyl-carnitines, analyte separation was achieved using an Agilent Poroshell 120, HILIC, 2.1×100 mm, 2.7 μm column. The pair of buffers used were 0.1% (v/v) formic acid and 10 mM ammonium formate in H_2_O and 0.1% (v/v) formic acid in acetonitrile, flowed at 200 μL/min on positive mode. MS1 data were acquired with the following settings: 3.5kV, capillarity temperature 300℃, 120 000, injection time 100ms, AGC 1×10^6^, scan range 200-1000. For ddMS2 data acquirement, the following settings were used: top 5, resolution 30,000, 200-2000, isolation 1.0m/z, nce 30, AGC target 1×10^3^, intensity threshold 5×10^3^, dynamic exclusion 20s. Raw data from both LC-MS/MS systems were extracted using MSConvert^69^ and in-house MATLAB scripts.

#### Quantifying ^13^C mass shifts from high-resolution MS1 and MS2 data

The extraction of mass spectrometry data was performed in MATLAB. First, retention times of CL, acyl carnitine, PC and PG species were confirmed using accurate precursor mass and MS2 fragmentation patterns. Accurate masses for the monoisotopic and expected mass shifts were calculated using ion formulas. The resulting ion cluster was used to visualise the analyte’s MS1 chromatogram and to integrate the area under the peak. These extracted MS1 ion intensities were then reported as ^13^C fractional enrichments^27^.

A labelled acyl chain could be incorporated into CLs with or without modification (e.g., elongation or desaturation). Thus, MS2 data was used to verify localisation of ^13^C-FA among the acyl chains. From the MS2 data we can derive the ratio of labelled to unlabelled product ions fragmented from a labelled precursor. Using relative ion intensities of labelled and unlabelled acyl anion [FA - H]^-^, the ion intensity of a labelled precursor (from MS1) is apportioned to its constituent acyl chains in terms of their labelled and unlabelled forms. The percentage labelling of a given acyl chain among CL or PG pools was then calculated by summing up these labelled and unlabelled constituents across all quantified CL or PG species.

#### Quantification and statistical analysis

Student’s t-test, one- or two-way ANOVA with Dunnett’s or Tukey’s multiple comparisons, Nonparametric Wilcoxon signed rank test, and linear regression statistical tests were performed with GraphPad Prism 9.0 (GraphPad Software). P < 0.05 was considered significant. Data are reported as mean ± standard error of the mean (SEM), standard deviation (SD), or min to max values of at least three independent determinations as indicated in figure legends. Schematic diagrams were created with Biorender.com.

## Acknowledgements

N.T.S. is supported by the Australian Rotary Health/Rotary Club of Blacktown City ‘Mel Grey’ PhD scholarship. L.M.B. and A.J.H. acknowledge grant support from The Movember Foundation/Prostate Cancer Foundation of Australia (MRTA3). A.J.H. is supported by a Robinson Fellowship from the University of Sydney and funding from the University of Sydney. L.M.B. is supported by a Principal Cancer Research Fellowship produced with the financial and other support of Cancer Council SA’s Beat Cancer Project on behalf of its donors and the State Government of South Australia through the Department of Health and was supported by a Future Fellowship from the Australian Research Council (FT130101004). We thank the Sydney Mass Spectrometry facility for access to LC-MS instruments; Julia Scott and Dr Zeyad Nassar in the Butler lab (University of Adelaide) for assistance with 3D spheroid and mitotracker experiment development; Dr Helen McGuire and the Sydney Flow Cytometry facility for assistance with mitotracker experiments; and Prof. Lisa Horvath and members of the Butler lab for scientific discussions.

## Author contributions

Conceptualisation by A.J.H. and L.E.Q. Project administration, visualisation, writing-original draft by N.T.S. Investigation by N.T.S., M.F.H.S., and A.W. Methodology by L.E.Q. Software by N.T.S. and L.E.Q. Formal analysis by N.T.S., L.E.Q and A.J.H. Writing-editing & reviewing by N.T.S., L.E.Q., A.J.H., and L.M.B. Funding acquisition, resources by A.J.H. Supervision by A.J.H., L.E.Q., and L.M.B.

## Extended Data

**Extended Data Fig. 1.**
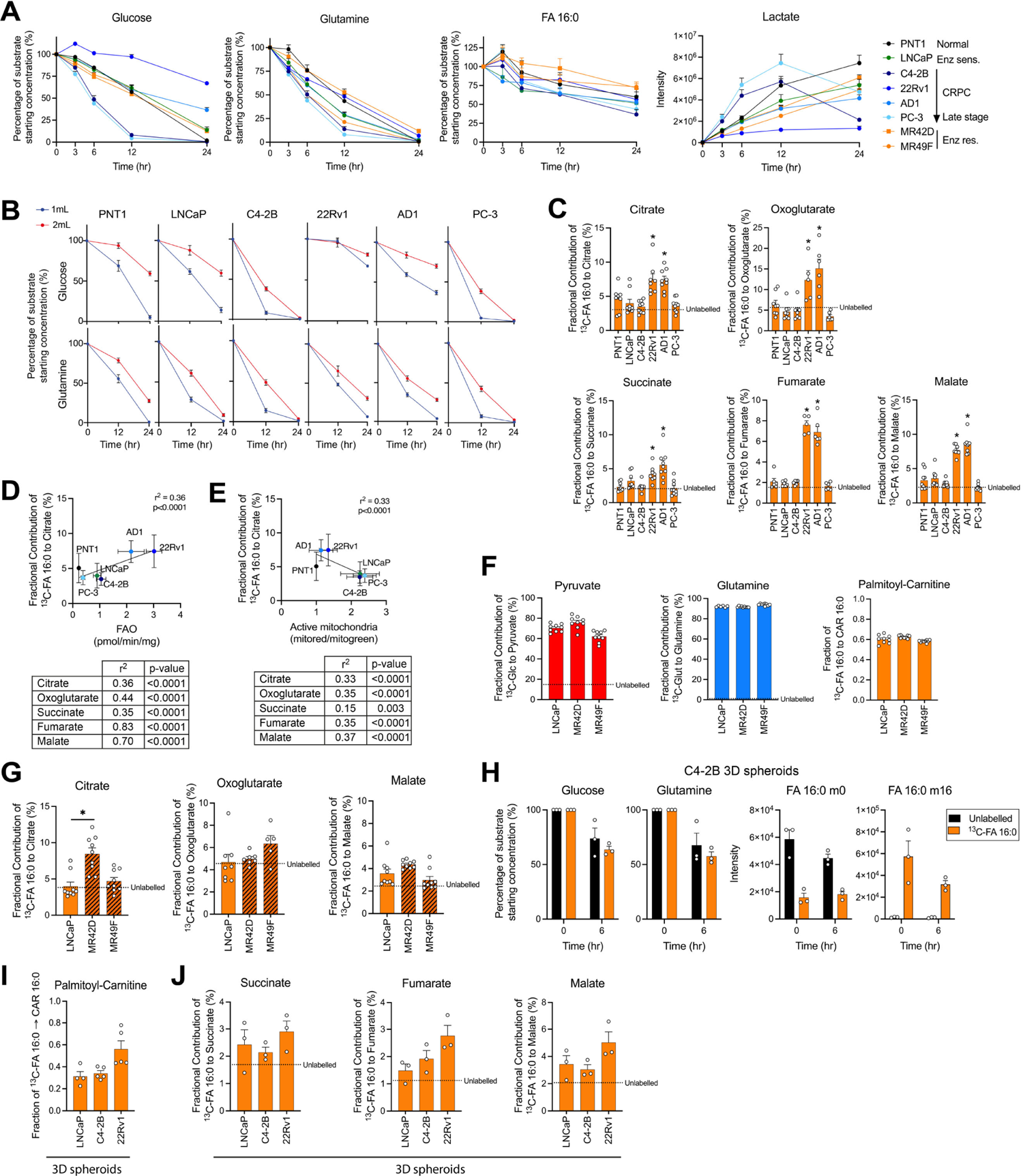
Palmitate is a minor carbon source to the TCA cycle. (A) Percentage of glucose, glutamine, and FA 16:0 relative to starting amount, and amount of lactate in culture media over 24 hours in 1mL culture medium. N=9 per cell line. (B) Percentage of glucose and glutamine relative to starting amount in 1 mL (blue) or 2 mL (red) culture medium. N=9 per cell line. (C) Fractional contribution of [U-^13^C]-FA16:0 to citrate, oxoglutarate, succinate, fumarate, and malate. N=9 per cell line. Statistically significant differences between PCa cells to PNT1 cells are shown. (D) Correlations of [U-^13^C]-FA 16:0 to citrate (plotted), oxoglutarate, succinate, fumarate, and malate to FAO. Error bars represent ± SD. P and r^2^ values by linear regression. (E) Correlations of [U-^13^C]-FA 16:0 to citrate (plotted), oxoglutarate, succinate, fumarate, and malate to active mitochondria content. Error bars represent ± SD. P and r^2^ values by linear regression. (F) Fractional contribution of [U-^13^C]-glucose to pyruvate (red), [U-^13^C]-glutamine to intracellular glutamine (blue), [U-^13^C]-FA 16:0 to palmitoyl-carnitine (orange) after 6 hours of labelling in LNCaP, MR42D and MR49F cells. N=9 per cell line. (G) Fractional contributions of [U-^13^C]-glucose, glutamine, or FA 16:0 to citrate, oxoglutarate and malate. N=9 per cell line. (H) Percentage of glucose, glutamine, and FA 16:0 m0 and m16 relative to starting amount, comparing unlabelled and [U-^13^C]-FA 16:0 media at 0 and 6 hours in representative C4-2B spheroids. N=3 per group. (I) Fractional contribution of [U-^13^C]-FA 16:0 to palmitoyl-carnitine in 3D spheroid LNCaP, C4-2B, and 22RV1 models. N=4-5 per cell line. (J) Fractional contribution of [U-^13^C]-FA 16:0 to succinate, fumarate, and malate in 3D spheroid models. N=3 per cell line. Graphs show mean ± SEM unless stated otherwise. *p<0.05 by One-way ANOVA with Dunnett’s multiple comparisons test, unless stated otherwise.

**Extended Data Fig. 2.**
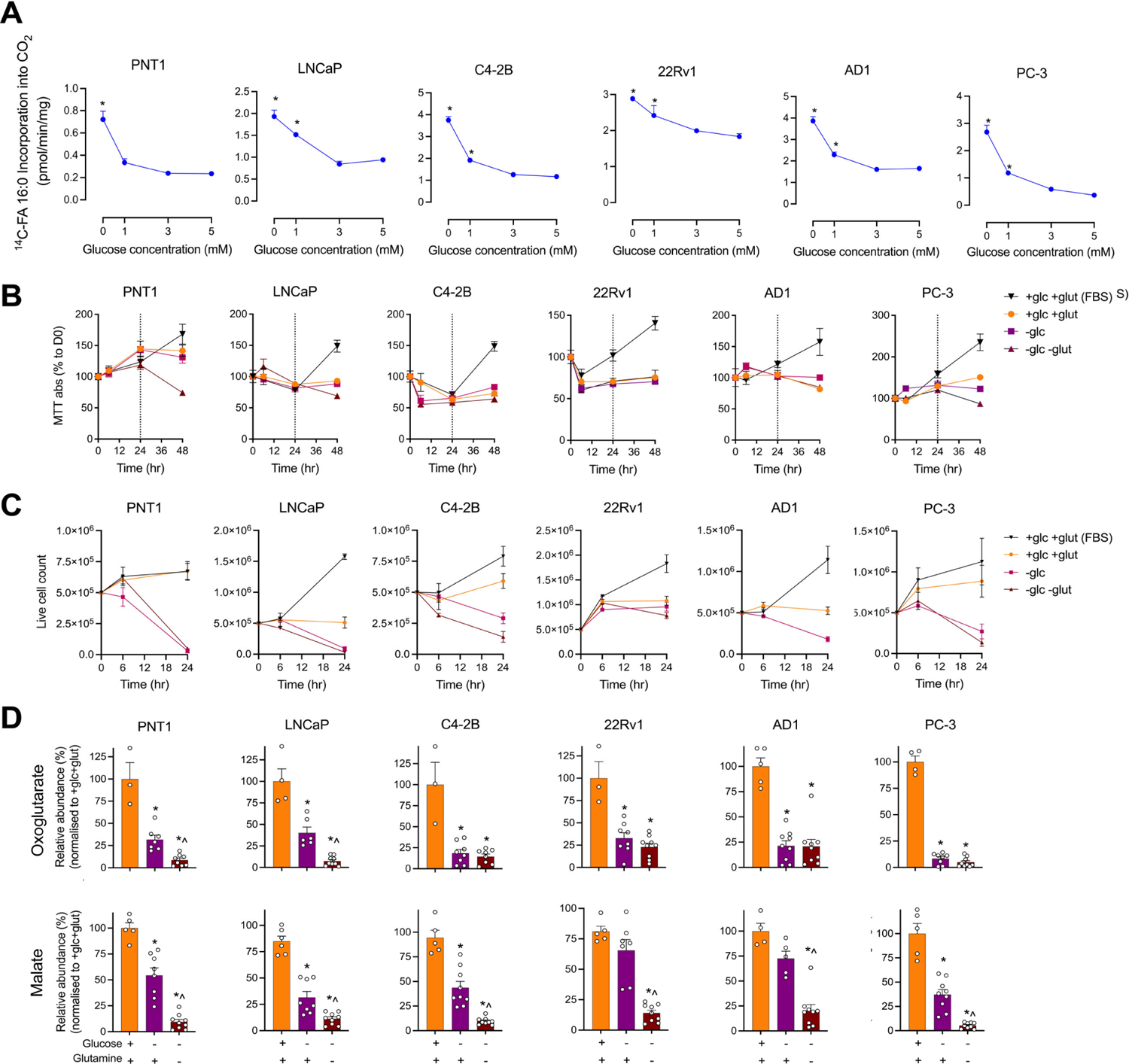
Palmitate cannot maintain TCA cycle activity under glucose and glutamine deprivation. (A) [1-^14^C]-FA 16:0 oxidation rates in the presence of 5 mM, 3 mM, 1 mM, or 0 mM glucose. N=3 per cell line. * compared to 5 mM glucose, p<0.05 by One-way ANOVA with Dunnett’s multiple comparisons test. (B) Impact of glucose- or glucose and glutamine-deprivation on metabolic activity assessed by MTT assay to 48 hours, normalised to day 0 results. Dotted line at 24 hours indicates replenishment of media. (C) Impact of glucose- or glucose and glutamine-deprivation on cell viability assessed by live cell counts at 0, 6 and 24 hours normalised to day 0 results. (D) Relative abundance (normalised to +glucose/+glutamine) of [U-^13^C]-FA 16:0 to oxoglutarate or malate. N=3-9 per cell line. Graphs show mean ± SEM. *compared to +glucose/+glutamine, ^compared to -glucose/+glutamine, p<0.05 by One-way ANOVA with Dunnett’s multiple comparisons test.

**Extended Data Fig. 3.**
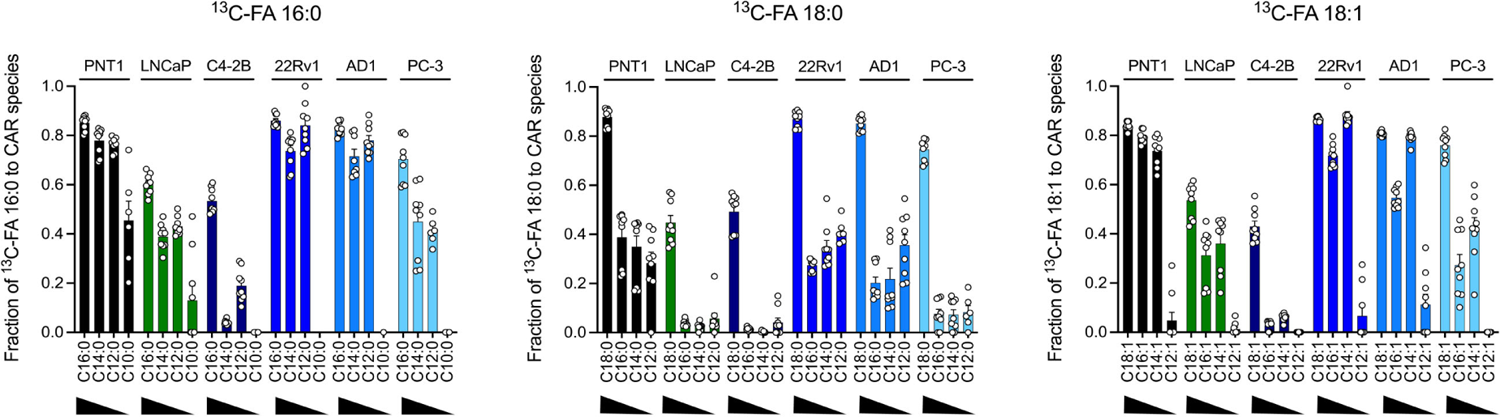
LCFAs undergo incomplete FAO in prostate cells. Fractional contributions of [U-^13^C]-FA 16:0, 18:0, 18:2 to shorter chain acyl-carnitine species. N=9 per cell line. Graphs show mean ± SEM.

**Extended Data Fig, 4.**
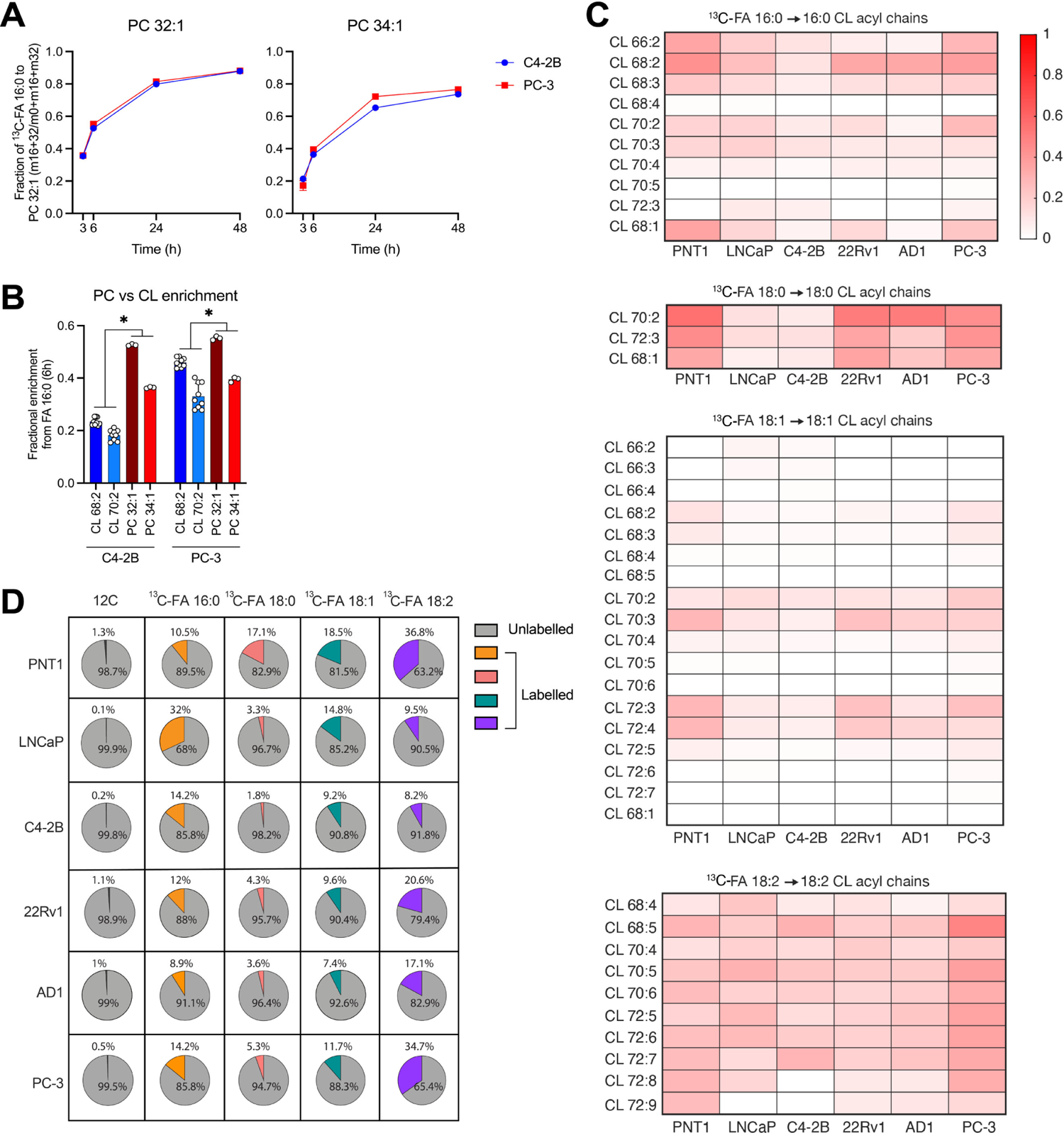
FA 18:2 predominantly enriches CL pools compared to other FAs. (A) Enrichment fraction of PC 32:1 and 34:1 in PC3 and C4-2B cells over 48 hours exposure to[U-^13^C]-FA 16:0. N=1 per cell line. (B) Comparison of fractional contribution of [U-^13^C]-FA 16:0 to PC and CL species at 6 hours in PC3 and C4-2B cells. N=3-9 per cell line. P<0.05 by two-way ANOVA with Tukey’s multiple comparisons test. (C) Fractional contribution of [U-^13^C]-FA 16:0, 18:0, 18:1, 18:2 (150 μM) to CL species containing the same acyl chains at 6 hours. N=3 per cell line. (D) Overall percentage of fatty acyls (of the same length and saturation as labelled substrate) contained in CL species labelled by [U-^13^C]-FA 16:0, 18:0, 18:1 or 18:2. Unlabelled ^12^C controls show level of background labelling. N=3 per cell line. Graphs show mean ± SEM unless stated otherwise.

**Extended Data Fig. 5.**
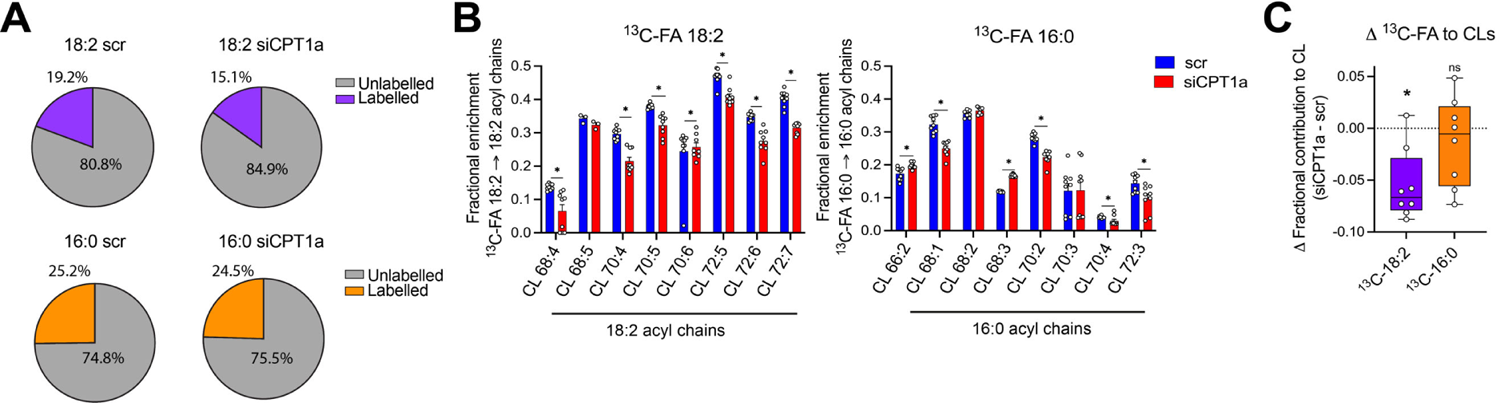
CPT1a^KD^ reduces FA 18:2 incorporation to CLs. (A) Percentage of 18:2 or 16:0 acyl chain in CLs respectively labelled by [U-^13^C]-FA 18:2 or 16:0 in scrambled (scr) and siCPT1a cells. N=9 per group. (B) Fractional contribution of [U-^13^C]-FA 18:2 or 16:0 to 18:2 or 16:0 acyl chains of CL species. N=9 per group. (C) Delta differences between fractional contribution of [U-^13^C]-FA 18:2 or 16:0 to acyl chains of CL species. N=7-8 per group. Statistically significant differences within [U-^13^C]-FA 18:2 or 16:0 groups to 0 difference, determined by nonparametric Wilcoxon signed rank test. Error bars represent ± min to max values. Graphs show mean ± SEM unless stated otherwise. ns, not significant, *p<0.05 by unpaired t-tests between siCPT1a and scr control groups unless stated otherwise.

## References

1 Swinnen, J. V., Esquenet, M., Goossens, K., Heyns, W. & Verhoeven, G. Androgens stimulate fatty acid synthase in the human prostate cancer cell line LNCaP. Cancer Res. 57, 1086–1090 (1997).

2 Yue, S. et al. Cholesteryl ester accumulation induced by PTEN loss and PI3K/AKT activation underlies human prostate cancer aggressiveness. Cell Metab. 19, 393–406, doi:10.1016/j.cmet.2014.01.019 (2014).

3 Migita, T. et al. Fatty acid synthase: a metabolic enzyme and candidate oncogene in prostate cancer. J. Natl. Cancer Inst. 101, 519–532, doi:10.1093/jnci/djp030 (2009).

4 Nassar, Z. D. et al. Human DECR1 is an androgen-repressed survival factor that regulates PUFA oxidation to protect prostate tumor cells from ferroptosis. Elife 9, doi:10.7554/eLife.54166 (2020).

5 Balaban, S. et al. Extracellular Fatty Acids Are the Major Contributor to Lipid Synthesis in Prostate Cancer. Mol. Cancer Res. 17, 949–962, doi:10.1158/1541-7786.Mcr-18-0347 (2019).

6 Fidelito, G. et al. Multi-substrate Metabolic Tracing Reveals Marked Heterogeneity and Dependency on Fatty Acid Metabolism in Human Prostate Cancer. Mol. Cancer Res. 21, 359–373, doi:10.1158/1541-7786.Mcr-22-0796 (2023).

7 Iglesias-Gato, D. et al. The Proteome of Primary Prostate Cancer. Eur. Urol. 69, 942–952, doi:10.1016/j.eururo.2015.10.053 (2016).

8 Butler, L. M. et al. Lipidomic Profiling of Clinical Prostate Cancer Reveals Targetable Alterations in Membrane Lipid Composition. Cancer Res. 81, 4981–4993, doi:10.1158/0008-5472.CAN-20-3863 (2021).

9 Ahmad, F., Cherukuri, M. K. & Choyke, P. L. Metabolic reprogramming in prostate cancer. Br. J. Cancer 125, 1185–1196, doi:10.1038/s41416-021-01435-5 (2021).

10 Costello, L. C. & Franklin, R. B. The intermediary metabolism of the prostate: a key to understanding the pathogenesis and progression of prostate malignancy. Oncology 59, 269–282, doi:10.1159/000012183 (2000).

11 Liu, Y. Fatty acid oxidation is a dominant bioenergetic pathway in prostate cancer. Prostate Cancer Prostatic Dis. 9, 230–234, doi:10.1038/sj.pcan.4500879 (2006).

12 Liu, Y., Zuckier, L. S. & Ghesani, N. V. Dominant uptake of fatty acid over glucose by prostate cells: a potential new diagnostic and therapeutic approach. Anticancer Res. 30, 369–374 (2010).

13 Van Heijster, F. H. A., Breukels, V., Jansen, K. F. J., Schalken, J. A. & Heerschap, A. Carbon sources and pathways for citrate secreted by human prostate cancer cells determined by NMR tracing and metabolic modeling. Proc. Natl. Acad. Sci. U. S. A. 119, doi:10.1073/pnas.2024357119 (2022).

14 Dueregger, A. et al. Differential Utilization of Dietary Fatty Acids in Benign and Malignant Cells of the Prostate. PLoS One 10, e0135704, doi:10.1371/journal.pone.0135704 (2015).

15 Mah, C. Y., Nassar, Z. D., Swinnen, J. V. & Butler, L. M. Lipogenic effects of androgen signaling in normal and malignant prostate. Asian J. Urol 7, 258–270, 10.1016/j.ajur.2019.12.003 (2020).

16 Fritz, I. B. & Yue, K. T. Long-chain carnitine acyltransferase and the role of acylcarnitine derivatives in the catalytic increase of fatty acid oxidation induced by carnitine. J. Lipid Res. 4, 279–288 (1963).

17 Carracedo, A., Cantley, L. C. & Pandolfi, P. P. Cancer metabolism: fatty acid oxidation in the limelight. Nat. Rev. Cancer 13, 227–232, doi:10.1038/nrc3483 (2013).

18 Ricciardi, M. R. et al. Targeting the leukemia cell metabolism by the CPT1a inhibition: functional preclinical effects in leukemias. Blood 126, 1925–1929, doi:10.1182/blood-2014-12-617498 (2015).

19 Schlaepfer, I. R. et al. Lipid Catabolism via CPT1 as a Therapeutic Target for Prostate Cancer. Mol. Cancer Ther. 13, 2361–2371, doi:10.1158/1535-7163.mct-14-0183 (2014).

20 Pike, L. S., Smift, A. L., Croteau, N. J., Ferrick, D. A. & Wu, M. Inhibition of fatty acid oxidation by etomoxir impairs NADPH production and increases reactive oxygen species resulting in ATP depletion and cell death in human glioblastoma cells. Biochim Biophys Acta 1807, 726–734, 10.1016/j.bbabio.2010.10.022 (2011).

21 Yao, C.-H. et al. Identifying off-target effects of etomoxir reveals that carnitine palmitoyltransferase I is essential for cancer cell proliferation independent of β-oxidation. PLoS Biol. 16, e2003782, doi:10.1371/journal.pbio.2003782 (2018).

22 O’Connor, R. S. et al. The CPT1a inhibitor, etomoxir induces severe oxidative stress at commonly used concentrations. Sci. Rep. 8, 6289, doi:10.1038/s41598-018-24676-6 (2018).

23 He, L. et al. Carnitine palmitoyltransferase-1b deficiency aggravates pressure overload-induced cardiac hypertrophy caused by lipotoxicity. Circulation 126, 1705–1716, doi:10.1161/circulationaha.111.075978 (2012).

24 Carta, G., Murru, E., Banni, S. & Manca, C. Palmitic Acid: Physiological Role, Metabolism and Nutritional Implications. Front. Physiol. 8, 902, doi:10.3389/fphys.2017.00902 (2017).

25 Ahn, W. S. & Antoniewicz, M. R. Parallel labeling experiments with [1,2-13C]glucose and [U-13C]glutamine provide new insights into CHO cell metabolism. Metab Eng. 15, 34–47, 10.1016/j.ymben.2012.10.001 (2013).

26 Duan, L. et al. 13C tracer analysis suggests extensive recycling of endogenous CO2 in vivo. Cancer Metab. 10, 11, doi:10.1186/s40170-022-00287-8 (2022).

27 Buescher, J. M. et al. A roadmap for interpreting (13)C metabolite labeling patterns from cells. Curr. Opin. Biotechnol. 34, 189–201, doi:10.1016/j.copbio.2015.02.003 (2015).

28 Bishop, J. L. et al. The Master Neural Transcription Factor BRN2 Is an Androgen Receptor–Suppressed Driver of Neuroendocrine Differentiation in Prostate Cancer. Cancer Discov. 7, 54–71, doi:10.1158/2159-8290.Cd-15-1263 (2017).

29 Joshi, M. et al. CPT1A Supports Castration-Resistant Prostate Cancer in Androgen-Deprived Conditions. Cells 8, 1115 (2019).

30 Nassar, Z. D. et al. Fatty Acid Oxidation Is an Adaptive Survival Pathway Induced in Prostate Tumors by HSP90 Inhibition. Mol. Cancer Res. 18, 1500–1511, doi:10.1158/1541-7786.mcr-20-0570 (2020).

31 Sutherland, R. M., McCredie, J. A. & Inch, W. R. Growth of Multicell Spheroids in Tissue Culture as a Model of Nodular Carcinomas2. J Natl Cancer Inst. 46, 113–120, doi:10.1093/jnci/46.1.113 (1971).

32 Jones, D. T. et al. 3D Growth of Cancer Cells Elicits Sensitivity to Kinase Inhibitors but Not Lipid Metabolism Modifiers. Mol. Cancer Ther. 18, 376–388, doi:10.1158/1535-7163.MCT-17-0857 (2019).

33 Tidwell, T. R., Røsland, G. V., Tronstad, K. J., Søreide, K. & Hagland, H. R. Metabolic flux analysis of 3D spheroids reveals significant differences in glucose metabolism from matched 2D cultures of colorectal cancer and pancreatic ductal adenocarcinoma cell lines. Cancer Metab. 10, 9, doi:10.1186/s40170-022-00285-w (2022).

34 Ma, Y. et al. Functional analysis of molecular and pharmacological modulators of mitochondrial fatty acid oxidation. Sci. Rep. 10, 1450, doi:10.1038/s41598-020-58334-7 (2020).

35 Sullivan, Lucas B. et al. Supporting Aspartate Biosynthesis Is an Essential Function of Respiration in Proliferating Cells. Cell 162, 552–563, 10.1016/j.cell.2015.07.017 (2015).

36 Le, A. et al. Glucose-Independent Glutamine Metabolism via TCA Cycling for Proliferation and Survival in B Cells. Cell Metab. 15, 110–121, doi:10.1016/j.cmet.2011.12.009 (2012).

37 Koves, T. R. et al. Peroxisome Proliferator-activated Receptor-γ Co-activator 1α-mediated Metabolic Remodeling of Skeletal Myocytes Mimics Exercise Training and Reverses Lipid-induced Mitochondrial Inefficiency. J. Biol. Chem. 280, 33588–33598, doi:10.1074/jbc.m507621200 (2005).

38 Koves, T. R. et al. Mitochondrial Overload and Incomplete Fatty Acid Oxidation Contribute to Skeletal Muscle Insulin Resistance. Cell Metab. 7, 45–56, doi:10.1016/j.cmet.2007.10.013 (2008).

39 Colbeau, A., Nachbaur, J. & Vignais, P. M. Enzymic characterization and lipid composition of rat liver subcellular membranes. Biochim Biophys Acta 249, 462–492, doi:10.1016/0005-2736(71)90123-4 (1971).

40 Falabella, M., Vernon, H. J., Hanna, M. G., Claypool, S. M. & Pitceathly, R. D. S. Cardiolipin, Mitochondria, and Neurological Disease. Trends Endocrinol Metab. 32, 224–237, doi:10.1016/j.tem.2021.01.006 (2021).

41 Ikon, N. & Ryan, R. O. Cardiolipin and mitochondrial cristae organization. Biochem Biophys Acta Biomembr. 1859, 1156–1163, 10.1016/j.bbamem.2017.03.013 (2017).

42 Schwall, C. T., Greenwood, V. L. & Alder, N. N. The stability and activity of respiratory Complex II is cardiolipin-dependent. Biochim Biophys Acta 1817, 1588–1596, 10.1016/j.bbabio.2012.04.015 (2012).

43 Dudek, J. et al. Cardiac-specific succinate dehydrogenase deficiency in Barth syndrome. EMBO Mol. Med. 8, 139–154, doi:10.15252/emmm.201505644 (2016).

44 Eble, K. S., Coleman, W. B., Hantgan, R. R. & Cunningham, C. C. Tightly associated cardiolipin in the bovine heart mitochondrial ATP synthase as analyzed by 31P nuclear magnetic resonance spectroscopy. J. Biol. Chem. 265, 19434–19440 (1990).

45 Taylor, W. A. & Hatch, G. M. Identification of the human mitochondrial linoleoyl-coenzyme A monolysocardiolipin acyltransferase (MLCL AT-1). J. Biol. Chem. 284, 30360–30371, doi:10.1074/jbc.M109.048322 (2009).

46 Zachman, D. K. et al. The role of calcium-independent phospholipase A2 in cardiolipin remodeling in the spontaneously hypertensive heart failure rat heart. J. Lipid Res. 51, 525–534, 10.1194/jlr.M000646 (2010).

47 Xu, Y. & Schlame, M. The turnover of glycerol and acyl moieties of cardiolipin. Chem. Phys. Lipids 179, 17–24, 10.1016/j.chemphyslip.2013.10.005 (2014).

48 Landriscina, C., Megli, F. M. & Quagliariello, E. Turnover of fatty acids in rat liver cardiolipin: Comparison with other mitochondrial phospholipids. Lipids 11, 61–66, doi:10.1007/BF02532585 (1976).

49 Wahjudi, P. N. et al. Turnover of nonessential fatty acids in cardiolipin from the rat heart. J. Lipid Res. 52, 2226–2233, doi:10.1194/jlr.M015966 (2011).

50 Nagarajan, S. R., Butler, L. M. & Hoy, A. J. The diversity and breadth of cancer cell fatty acid metabolism. Cancer Metab 9, 2, doi:10.1186/s40170-020-00237-2 (2021).

51 Oemer, G. et al. Fatty acyl availability modulates cardiolipin composition and alters mitochondrial function in HeLa cells. J. Lipid Res. 62, 100111, doi:10.1016/j.jlr.2021.100111 (2021).

52 Hoch, F. L. Cardiolipins and biomembrane function. Biochimica et Biophysica Acta (BBA) 1113, 71–133, doi:10.1016/0304-4157(92)90035-9 (1992).

53 Cao, J., Liu, Y., Lockwood, J., Burn, P. & Shi, Y. A Novel Cardiolipin-remodeling Pathway Revealed by a Gene Encoding an Endoplasmic Reticulum-associated Acyl-CoA:Lysocardiolipin Acyltransferase (ALCAT1) in Mouse. J. Biol. Chem. 279, 31727–31734, doi:10.1074/jbc.m402930200 (2004).

54 Lu, Y.-W. & Claypool, S. M. Disorders of phospholipid metabolism: an emerging class of mitochondrial disease due to defects in nuclear genes. Frontiers in Genetics 6, doi:10.3389/fgene.2015.00003 (2015).

55 Xu, Y., Malhotra, A., Ren, M. & Schlame, M. The Enzymatic Function of Tafazzin*. J. Biol. Chem. 281, 39217–39224, 10.1074/jbc.M606100200 (2006).

56 Hostetler, K. Y., Galesloot, J. M., Boer, P. & Van Den Bosch, H. Further studies on the formation of cardiolipin and phosphatidylglycerol in rat liver mitochondria. Effect of divalent cations and the fatty acid composition of CDP-diglyceride. Biochim Biophys Acta 380, 382–389, doi:10.1016/0005-2760(75)90106-x (1975).

57 Zadra, G. et al. Inhibition of de novo lipogenesis targets androgen receptor signaling in castration-resistant prostate cancer. Proc. Natl. Acad. Sci. U. S. A. 116, 631–640, doi:10.1073/pnas.1808834116 (2019).

58 Parida, P. K. et al. Limiting mitochondrial plasticity by targeting DRP1 induces metabolic reprogramming and reduces breast cancer brain metastases. Nature Cancer 4, 893–907, doi:10.1038/s43018-023-00563-6 (2023).

59 Yang, J. H. et al. Snail augments fatty acid oxidation by suppression of mitochondrial ACC2 during cancer progression. Life Science Alliance 3, e202000683, doi:10.26508/lsa.202000683 (2020).

60 Flaig, T. W. et al. Lipid catabolism inhibition sensitizes prostate cancer cells to antiandrogen blockade. Oncotarget 8, 56051–56065, doi:10.18632/oncotarget.17359 (2017).

61 Zhang, H. et al. Lipidomics reveals carnitine palmitoyltransferase 1C protects cancer cells from lipotoxicity and senescence. J Pharm Anal 11, 340–350, doi:10.1016/j.jpha.2020.04.004 (2021).

62 Paradies, G., Paradies, V., De Benedictis, V., Ruggiero, F. M. & Petrosillo, G. Functional role of cardiolipin in mitochondrial bioenergetics. Biochim Biophys Acta 1837, 408–417, 10.1016/j.bbabio.2013.10.006 (2014).

63 Zhong, H. et al. Mitochondrial control of apoptosis through modulation of cardiolipin oxidation in hepatocellular carcinoma: A novel link between oxidative stress and cancer. Free Radic. Biol. Med. 102, 67–76, 10.1016/j.freeradbiomed.2016.10.494 (2017).

64 Faubert, B. et al. Lactate Metabolism in Human Lung Tumors. Cell 171, 358–371.e359, doi:10.1016/j.cell.2017.09.019 (2017).

65 Neinast, M. D. et al. Quantitative Analysis of the Whole-Body Metabolic Fate of Branched-Chain Amino Acids. Cell Metab. 29, 417–429.e414, 10.1016/j.cmet.2018.10.013 (2019).

66 Kiebish, M. A., Han, X., Cheng, H., Chuang, J. H. & Seyfried, T. N. Cardiolipin and electron transport chain abnormalities in mouse brain tumor mitochondria: lipidomic evidence supporting the Warburg theory of cancer. J. Lipid Res. 49, 2545–2556, doi:10.1194/jlr.M800319-JLR200 (2008).

67 Nyquist, M. D. et al. TALEN-engineered AR gene rearrangements reveal endocrine uncoupling of androgen receptor in prostate cancer. Proc. Natl. Acad. Sci. U. S. A. 110, 17492–17497, doi:doi:10.1073/pnas.1308587110 (2013).

68 Li, Y. et al. Androgen receptor splice variants mediate enzalutamide resistance in castration-resistant prostate cancer cell lines. Cancer Res. 73, 483–489, doi:10.1158/0008-5472.Can-12-3630 (2013).

69 Chambers, M. C. et al. A cross-platform toolkit for mass spectrometry and proteomics. Nat. Biotechnol. 30, 918–920, doi:10.1038/nbt.2377 (2012).

70 Lu, W. et al. Metabolite Measurement: Pitfalls to Avoid and Practices to Follow. Annu. Rev. Biochem. 86, 277–304, doi:10.1146/annurev-biochem-061516-044952 (2017).

